# Allosteric inhibition of the epidermal growth factor receptor through disruption of transmembrane interactions

**DOI:** 10.1101/2022.10.31.514582

**Authors:** Jennifer A Rybak, Amita R Sahoo, Soyeon Kim, Robert J Pyron, Savannah B Pitts, Saffet Guleryuz, Adam W Smith, Matthias Buck, Francisco N Barrera

## Abstract

The epidermal growth factor receptor (EGFR) is a receptor tyrosine kinase (RTK) commonly targeted for inhibition by anti-cancer therapeutics. Current therapeutics target EGFR’s kinase domain or extracellular region. However, these types of inhibitors are not specific for tumors over healthy tissue and therefore cause undesirable side effects. Our lab has recently developed a new strategy to regulate RTK activity by designing a peptide that specifically binds to the transmembrane (TM) region of the RTK to allosterically modify kinase activity. These peptides are acidity-responsive, allowing them to preferentially target acidic environments like tumors. We have applied this strategy to EGFR and created the PET1 peptide. We observed that PET1 behaves as a pH-responsive peptide that modulates the configuration of the EGFR TM through a direct interaction. Our data indicated that PET1 inhibits EGFR-mediated cell migration. Finally, we investigated the mechanism of inhibition through molecular dynamics simulations, which showed that PET1 sits between the two EGFR TM helices; this molecular mechanism was additionally supported by AlphaFold-Multimer predictions. We propose that the PET1-induced disruption of native TM interactions disturbs the conformation of the kinase domain in such a way that it inhibits EGFR’s ability to send migratory cell signals. This study is a proof-of-concept that acidity-responsive membrane peptide ligands can be generally applied to RTKs. In addition, PET1 constitutes a viable approach to therapeutically target the TM of EGFR.

## Introduction

The epidermal growth factor receptor (EGFR) is a HER-family receptor tyrosine kinase (RTK) that is involved in cell signaling in healthy tissue. Activation of EGFR regulates essential cellular processes including cell migration, proliferation, and apoptosis (1). To mediate these processes, the extracellular ligand binding region of EGFR senses environmental cues via interactions with one of its 7 known ligands, of which epidermal growth factor (EGF) is the most well characterized (2, 3). Ligand binding promotes EGFR oligomerization mediated by the extracellular region. Signaling is then transduced across the membrane by altering the configuration of the transmembrane (TM) domain, by dimerization of the TM helical region or a change in the arrangement of the TM helices within such a dimer. Specifically, the TM of unliganded (inactive) EGFR dimerizes at the C-terminus (C_t_), while the ligand bound form dimerizes N-terminally (N_t_), and the two helices are also rotated by 180° between the conformations (4, 5). The ligand-bound TM configuration promotes asymmetric dimerization of the intracellular juxta-membrane (JM) and kinase domains, which causes autophosphorylation of intracellular tyrosine residues (6, 7). Effector proteins are then recruited to activate various cellular signaling pathways, including RAS/RAF/MEK, PI3K/AKT/mTOR, and JAK/STAT (1).

Because of its essential roles in cell signaling, misregulation or overexpression of EGFR often causes a cancerous phenotype. Indeed, EGFR is commonly overexpressed in solid tumors, such as breast, colon, head-and-neck, renal, ovarian, and non–small-cell lung cancer (8–11). Furthermore, EGFR-mediated cancers tend to be more aggressive (12). Currently, there are two main therapeutic approaches that are effective for targeting EGFR in cancer: monoclonal antibodies and small-molecule tyrosine kinase inhibitors (TKIs) (11). Both approaches are generally safer and more efficacious than chemotherapy. However, major challenges for antibodies include a short lifespan and variations in tumor development (13), and TKIs are often promiscuous amongst other RTKs due to the highly conserved ATP binding pocket causing off-target effects (11, 14). Additionally, both strategies often become less effective over time as tumors develop resistance (11, 13, 14). For these reasons, it is necessary to find safer, more effective, and more selective ways to inhibit EGFR activity.

Recently, our group has developed a novel approach to modify RTK activity by targeting the receptor’s TM domain using a pH-responsive peptide (15). Acidity responsive peptides such as pHLIP (16) and ATRAM (17, 18) are marginally hydrophobic and contain acidic residues across the sequence. At physiological pH, the acidic residues are unprotonated and therefore negatively charged, allowing the peptide to be soluble, but able to bind to the surface of lipid membranes, in an unstructured conformation. At lower pH, the acidic residues become protonated, resulting in membrane insertion and a gain of α-helical structure. These peptides can preferentially target cancer cells over healthy tissue by taking advantage of the slightly acidic extracellular pH that is hallmark of tumors (18–22). Alves et al. (15) evolved this concept and designed the peptide TYPE7 to be specific for the RTK EphA2. TYPE7 binds the TM to allosterically regulate EphA2 kinase activity by causing a configurational change (23). This method of regulation is likely to be more selective for EphA2 than targeting the highly conserved kinase domain. When combined with the increased selectivity for cancer cells, TYPE7 represents a potentially useful development to target RTK activity in cancer.

We sought to use the TYPE7 approach to inhibit EGFR. Here we report a novel pH-responsive <u>P</u>eptide for <u>E</u>GFR <u>T</u>argeting (PET1). PET1 binds selectively to EGFR in cancer cells and inhibits the ability of the ligand EGF to activate cell migration. Interestingly, PET1 does not modify the receptor’s oligomerization state. Molecular dynamics simulations and AlphaFold-Multimer predictions reveal that PET1 disrupts EGFR by forcing apart the TM dimer, bridging the individual TM domains. PET1 therefore induces a configuration of EGFR that is inactive without full dissociation of the oligomeric complex.

## Results

### PET1 is a pH-responsive peptide that interacts with the TM region of EGFR

To design PET1, we modified the transmembrane region of human EGFR as previously described (Figure 1A) (15). Glutamic acid (E) residues were strategically placed throughout the TM region and at the charged JM region immediately C_t_ to the TM. Glutamic acid residues are negatively charged at neutral pH and only become protonated at low pH, and therefore are expected to confer pH-responsiveness to the peptide. To determine the interaction of PET1 with lipid membranes, we performed complementary biophysical assays. We used circular dichroism (CD) and oriented circular dichroism (OCD) to assess PET1 secondary structure and the average tilt of membrane insertion, respectively (24). CD performed in buffer (Figure 1B, black) and POPC lipid vesicles at pH 7.5 (Figure 1B, grey) revealed PET1 is in a random coil conformation in both conditions, as indicated by the minima at 200 nm. However, in vesicles at pH 4.2, PET1 adopts an α-helical conformation based on the minima at 208 and 222 nm (Figure 1B, red), indicating that PET1 is only structured at acidic pH. To determine if folding is due to membrane insertion, we performed OCD at acidic pH. OCD measures the average tilt relative to the membrane normal of an α-helix. The OCD curve of PET1 (Figure 1C) revealed a peptide configuration that is inserted but with a noticeable helical tilt, as indicated by the similar intensity at 208 and 225 nm. Such orientation is consistent with NMR structures of the EGFR TM domain (4, 25, 26). Together the biophysical results reveal that PET1 is pH-responsive because the peptide undergoes the desired shift from unstructured to TM the pH decreases.

**Figure 1:**
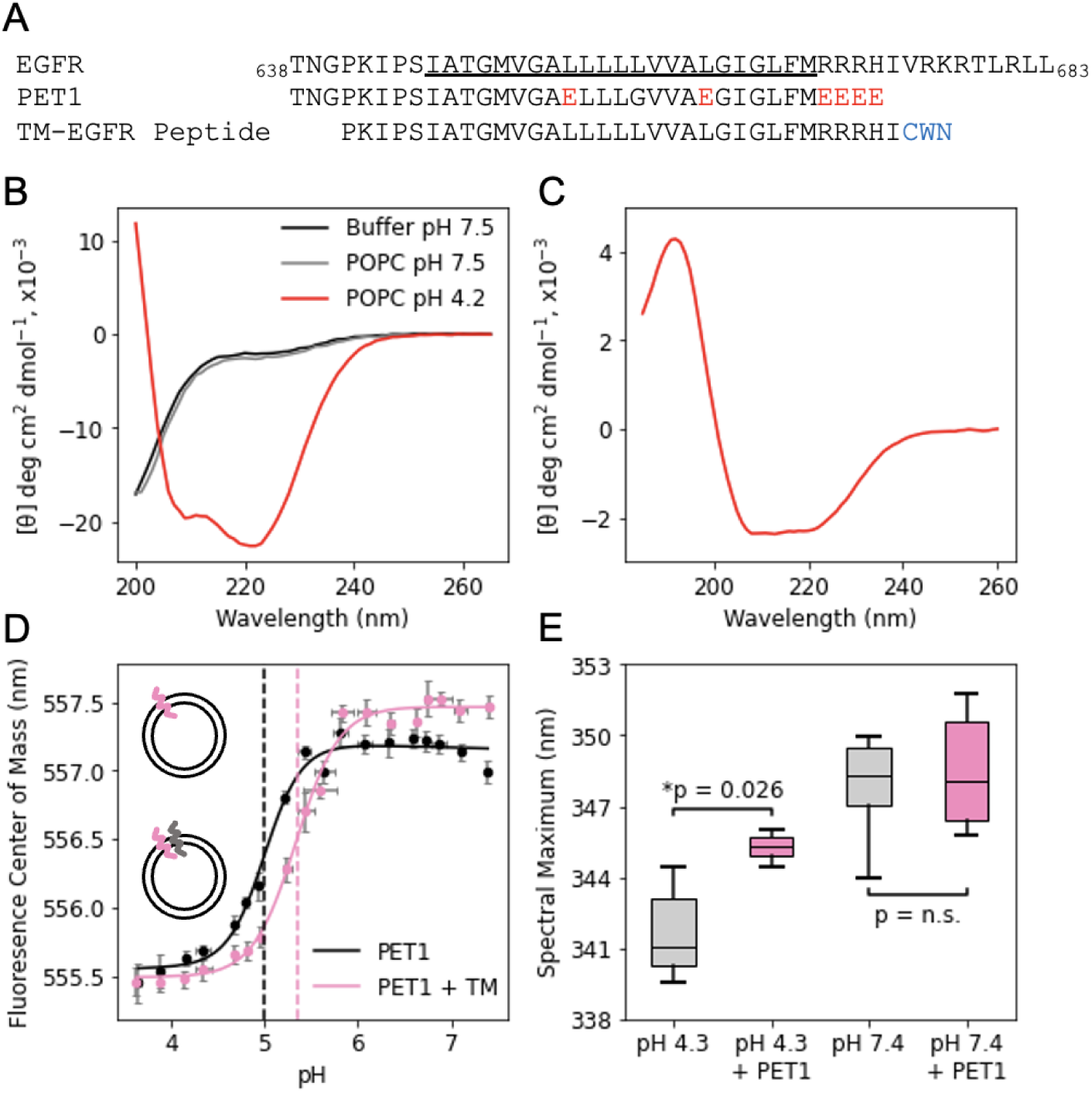
PET1 is a pH-responsive transmembrane peptide that binds to the TM of EGFR. **A,** The sequence of EGFR (residue 638 to 683) with the TM domain underlined is aligned with the sequence of PET1 and TM-EGFR peptides. Amino acid mutations to acidic residues are highlighted in red. The C_t_ CWN tag on TM-EGFR is highlighted in blue. **B,** CD spectra informs on the secondary structure changes of PET1 in buffer at pH 7.5 (black), in POPC vesicles at pH 7.5 (grey), and in POPC vesicles at pH 4.2 (red). **C,** Oriented circular dichroism of PET1 in POPC supported bilayers at pH 4.2 (red). **D,** The fluorescence center of mass of PET1-NBD in POPC vesicles alone (black, top cartoon) or containing TM-EGFR (pink, bottom cartoon) was determined at varying pH values to determine the pH_50_ of insertion (dashed lines) using Equation 2. Reported pH_50_ values (mean ± S.D.) are an average of 3 individual replicates. Statistical analysis was performed using a *t*-test (*p* = 0.029). **E,** The fluorescence spectra was recorded for the C_t_ W residue of TM-EGFR in POPC vesicles at pH 4.3 and 7.4 in the presence (pink) or absence (grey) of PET1. Box plot conveys the wavelength corresponding to the maximum fluorescence of the curve. N = 6. Statistical analysis was performed using a One-Way ANOVA (*p* = 3 x 10^-5^) with a post-hoc Dunnet-T3 for comparisons between groups, as the Levene Statistic was significant (*p* = 0.015).

We performed a pH titration experiment to further investigate the pH-dependent membrane insertion of PET1. A fluorescently labeled PET1 (PET1-NBD, Supp. Figure 1) was incubated with POPC vesicles, and the pH was changed to cover a wide pH range from acidic to neutral values. NBD is an environmentally sensitive dye that presents a blue-shifted fluorescence spectra in a hydrophobic environment (27–29). Therefore, we used the NBD fluorescence center of mass (Equation 1) as an indicator for the peptide environment (Figure 1D). At low pH, we observed a low center of mass that suggests a more hydrophobic (probably membrane-associated) environment, while the higher center of mass at neutral pH suggests that the NBD in PET1 is more solvent-exposed. The transition between states occurred in a sigmoidal fashion with a midpoint (pH_50_) of 5.00 ± 0.01 (24). For pH-responsive peptides that bind the TM of an RTK, the presence of that RTK can increase the pH_50_ due to an increase in tendency to be inserted in the presence of a membrane binding partner (15, 23). For this reason, we repeated the experiment using proteo-liposomes containing a peptide mimic of the TM of EGFR (TM-EGFR). Under these conditions, we observed that the pH_50_ value increased to 5.36 ± 0.16 (Figure 1D). This result indicates that the presence of TM-EGFR makes PET1 insertion more favorable, suggesting that PET1 interacts with the TM region of EGFR.

PET1-NBD experiments described the effect that TM-EGFR causes in PET1 insertion. We performed an orthogonal experiment to determine if PET1 also affects the configuration of TM-EGFR. For this, we used the C_t_ tryptophan (W) residue on TM-EGFR as a fluorescent reporter of hydrophobicity, similarly to NBD. W also presents a blue shift in more hydrophobic environments (30). Using pH values representative of the fully TM (pH 4.3) or fully unstructured (pH 7.4) PET1 baselines as determined by the titration experiment, we measured the W fluorescence spectra of TM-EGFR in POPC lipid vesicles with and without PET1 added. We used the spectral maximum as an indicator for W positioning. At pH 4.3 when PET1 is fully inserted, the addition of PET1 caused a significant increase in the spectral max wavelength (Figure 1E). This wavelength increase was accompanied by a significant fluorescence decrease (Supp. Figure 2). The observed spectral red-shift indicates a transition to a more polar environment when PET1 forms a TM helix. At pH 7.4 when PET1 does not insert to the membrane, the addition of PET1 had no effect (Figure 1D, Supp. Figure 2). These results suggest that only the TM conformation of PET1 modifies the environment of the TM-EGFR C_t_, due to an interaction between PET1 and TM-EGFR.

**Figure 2:**
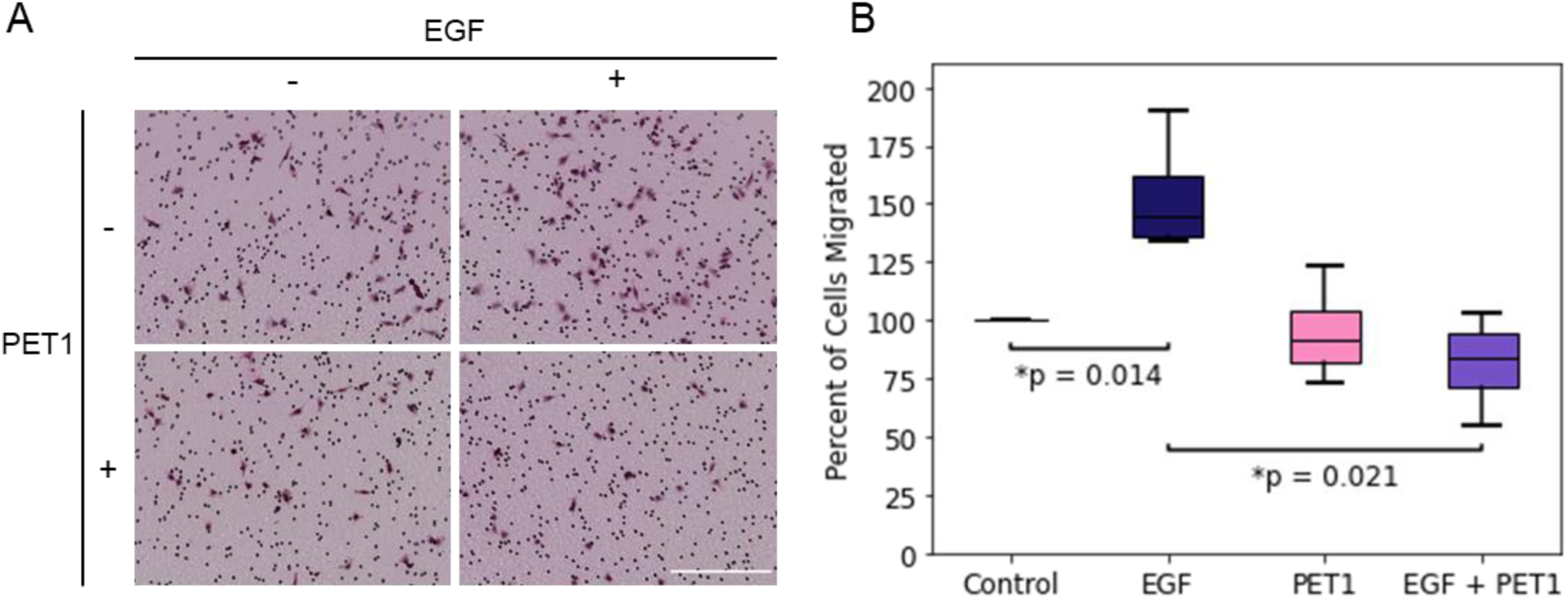
PET1 inhibits the migratory response to EGF. **A,** Boyden chamber assay performed in A375 cells with no treatment, or with EGF (100 ng/mL), PET1 (2 µM), and EGF + PET1. Representative images are shown. The small black dots are membrane pores and cells are stained purple. Scale bar = 250 µm. **B,** Box plot shows compiled migration data. N = 3 with each biological replicate normalized to Control conditions. Statistical analysis was performed using a Kruskal Wallis test -H(3) = 9.761, *p* = 0.021-with a Mann Whitney U test for comparisons between groups.

### PET1 inhibits EGFR-mediated cell migration

We next sought to determine if PET1 modifies EGFR activity. We used cell migration as an indicator of downstream EGFR-regulated cell signaling, since activation of this RTK promotes cell migration. We performed a Boyden cell chamber assay in which A375 melanoma cells migrate through a porous membrane in response to a chemoattractant (Figure 2). As expected, treatment with EGF significantly enhanced cell migration due to its ability to activate EGFR (31, 32). We observed that PET1 alone did not change basal levels of migration, but interestingly, PET1 was able to significantly block the ability of EGF to promote migration. We have previously shown that the pHLIP peptide, of similar pH-responsive properties as PET1, is not able to affect cell migration of A375 cells (33), which suggests that effect of PET1 is specific to its interaction with EGFR. To validate the migration results, we performed a control cell viability assay using the MTS reagent (Supp. Figure 3). We found that PET1 caused no significant cellular toxicity, with or without EGF treatment. Our data indicate that PET1 inhibits ligand-induced activation of EGFR.

**Figure 3:**
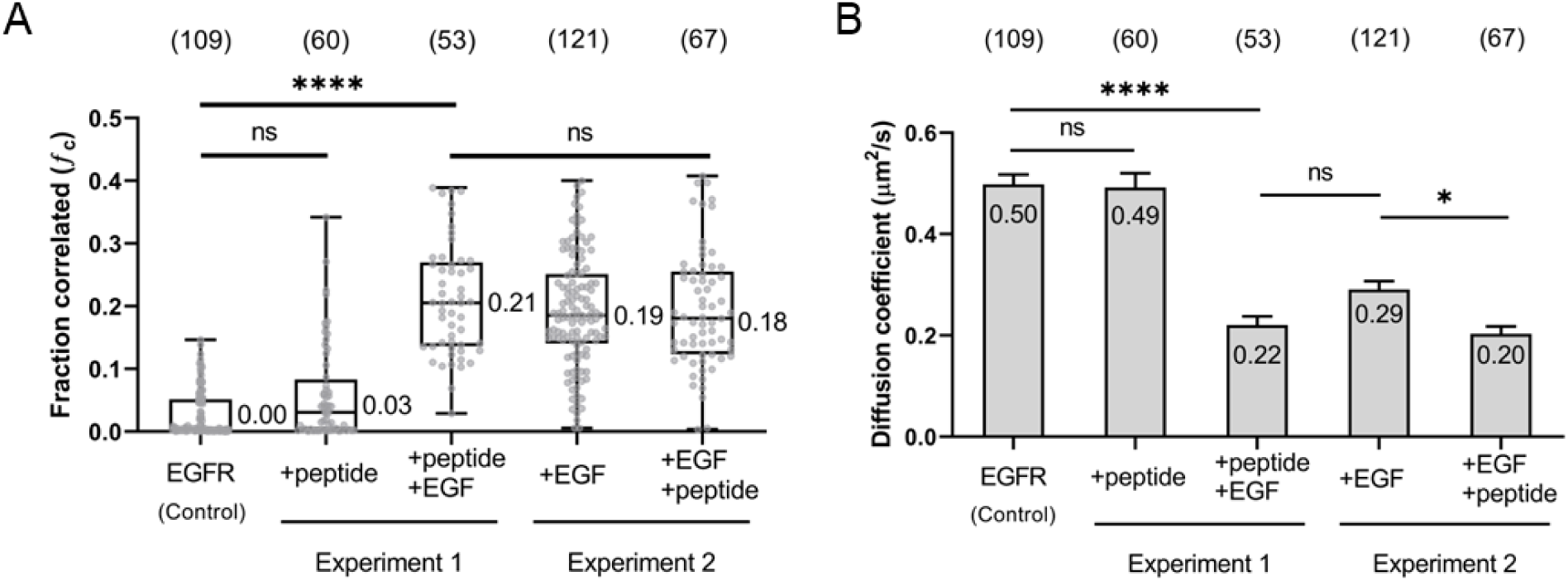
PET1 does not affect the oligomerization state of EGFR. **A,** Cross correlation values of EGFR in the presence of PET1 peptide or ligand (EGF) stimulation. Each data point is a single cell (Cos7) measurement (total number shown at top in parenthesis). Box and whisker plots were generated to visualize the 25-75 percentile and median values of the distributions. In Experiment 1, we added EGF after PET1 treatment, and in Experiment 2 we added PET1 after EGF treatment. **B,** Diffusion coefficient values from the same single cell measurements summarized in the left panel. The height of the bars is the mean and the error bars are the standard error (SEM).

### PET1 does not alter EGFR oligomerization

We investigated next the mechanism through which PET1 inhibits EGFR activation. Activation of EGFR by EGF promotes receptor self-assembly by stabilizing the dimeric and oligomeric states. We therefore studied the effect of PET1 on EGFR oligomerization via pulsed interleaved excitation fluorescence cross-correlation spectroscopy (PIE-FCCS) (34). PIE-FCCS is a time-resolved fluorescence method in which two excitation lasers are focused on the plasma membrane of live cells and fluorescence fluctuations are recorded to quantify the expression level, mobility, and oligomerization state of the labeled membrane proteins. Single cell data are fit to determine the fraction of the co-diffusing species relative to the red or green species (*f_c_*). We assessed the oligomerization state of EGFR before and after PET1 addition by comparing *f_c_* values (Figure 3A). In order to determine the effect of the peptide on EGFR receptor-receptor interactions, we performed two sets of experiments. In the first, we collected data after PET1 was added to EGFR-expressing cells in the absence of ligand. Then, we added EGF to determine if the peptide affected ligand-stimulated multimerization of EGFR. In unstimulated cells, the median *f_c_* value was 0.00, indicating that EGFR was predominantly monomeric. Upon peptide addition, the median *f_c_* value was 0.03, with no statistical difference compared to unstimulated EGFR. Upon EGF addition to these PET1 treated cells, a median *f_c_* value of 0.21 was obtained. This value is significantly larger, confirming that EGF considerably promotes ligand-stimulated EGFR multimerization even in the presence of PET1.

The second set of the experiments was to test if a multimeric state, generated by first adding EGF ligand to EGFR, could be disrupted by PET1. First, we collected data after EGF was added to EGFR-expressing cells. We then added PET1 to the media and collected more PIE-FCCS measurements. EGF-stimulated EGFR yielded a median *f_c_* value of 0.19. When the peptide was added to the well, the median *ƒ_c_* values was 0.18. There was no significant difference between the *f_c_* values (Figure 3A) of EGFR oligomers, regardless of the presence of the peptide, indicating that PET1 does not disrupt the oligomerization of EGFR. In fact, the diffusion of the EGFR:PET1 complex is slightly slower than the diffusion of the EGFR oligomer alone (Figure 3B).

### PET1 co-localizes with and binds to EGFR

To demonstrate that the effect of PET1 was direct, we sought to demonstrate the interaction between PET1 and EGFR. We first determined the cellular location of PET1 with respect to EGFR by treating A431 cells, which have a naturally high level of EGFR expression, with a fluorescently tagged version of PET1 (PET1-DL680, Supp. Figure 1). We visualized the cellular location of PET1 and EGFR via confocal microscopy (Figure 4A). PET1 localized at the plasma membrane, and there was a strong overlap between PET1 and EGFR signals irrespective of EGF treatment. To quantify co-localization, we determined the Pearson’s correlation coefficient (*r*) (Figure 4B). The *r* parameter ranges from +1 for perfect correlation to −1 for anti-correlation (35). *r* was 0.7 with and without EGF, indicating a strong correlation between PET1-DL680 and EGFR cellular localization regardless of whether EGF is present. This result suggests that PET1 can bind to EGFR prior to engagement with EGF.

**Figure 4:**
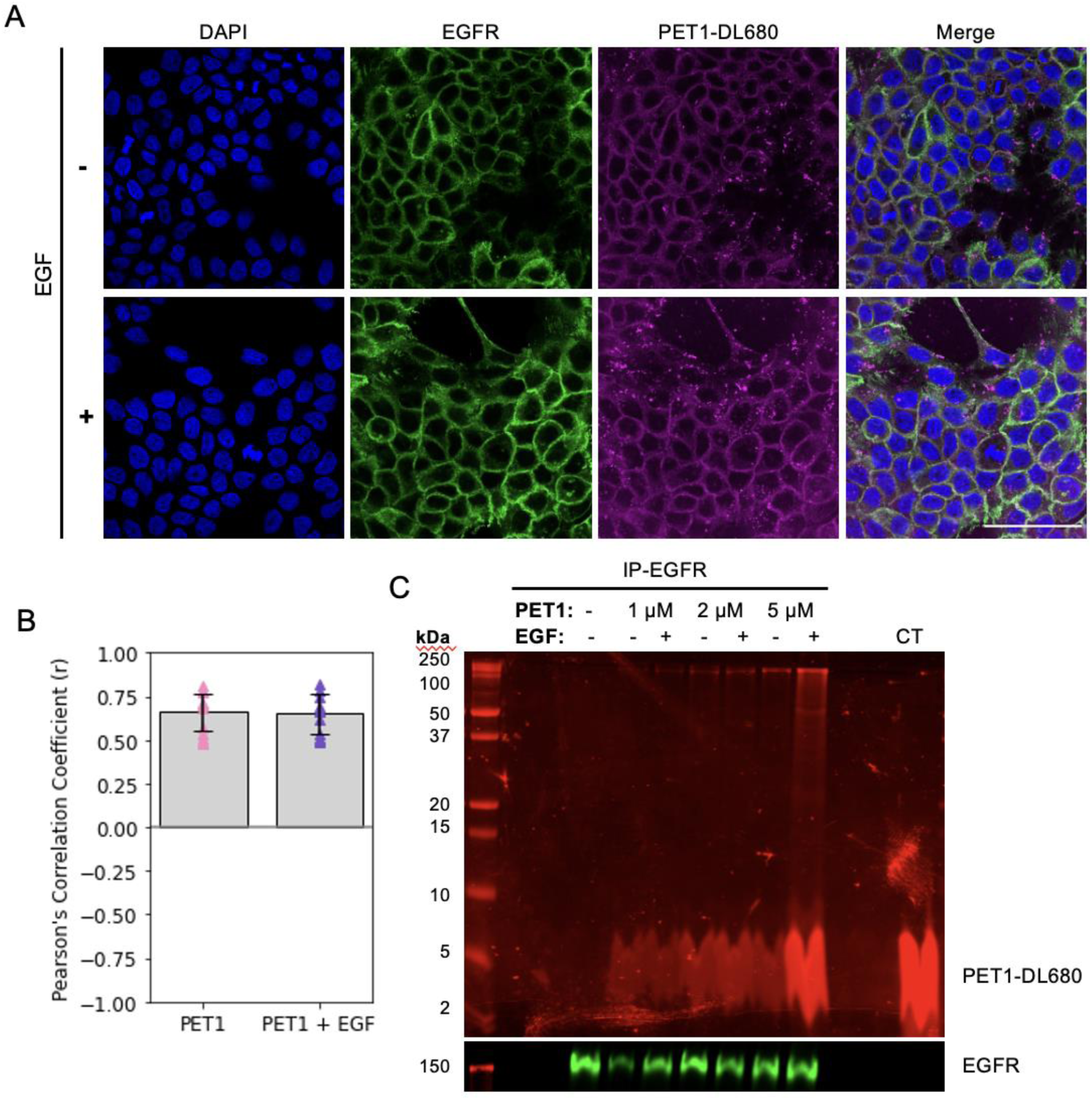
PET1 interacts with endogenous EGFR. **A,** Representative images of A431 cells treated with PET1-DL680 (magenta) for one hour followed by a 5 min incubation with or without EGF, fixed and stained for EGFR (green). DAPI was used for nucleus staining (blue).. Scale = 75 µm. **B,** Pearson’s correlation coefficient (*r*) was calculated for PET1 and EGFR channels. N = 3, n = 15. The error bars represent standard deviation of the mean. **C,** EGFR was immunoprecipitated from lysates of A431 cells treated with PET1-DL680 for one hour followed by 5 min treatment with or without EGF. Control (CT) lane shows that PET1-DL680 alone runs as a wide band. SDS-PAGE and Western blot of the eluates for EGFR (green) is shown.

Next, we performed co-immunoprecipitation (co-IP) experiments to investigate whether co-localization was due to EGFR-PET1 binding. We immunoprecipitated EGFR from cells treated with PET1-DL680 using the gentle detergent NP-40 to allow for precipitation of bound proteins. Figure 4C shows that PET1-DL680 at varying concentrations co-IP’s with EGFR. We also observed that binding is independent of the presence of EGF, in agreement with the co-localization results. In addition to the low molecular weight band corresponding to PET1-DL680 (broad band at ∼4 kDa), we observed a high molecular weight fluorescent band likely due to an SDS-resistant complex of PET1-DL680 and EGFR (∼180 kDa). Co-immunoprecipitation and SDS-resistant binding suggest that the interaction between PET1-DL680 and EGFR is strong.

To determine if PET1 action on EGFR is selective, we utilized an array of 58 RTKs to measure phosphorylation in control conditions and in the presence of EGF. We studied the effect of PET1 and used a scrambled peptide as a negative control (Supp. Figure 4). PET1 alone did not cause phosphorylation of any RTKs, suggesting that the peptide does not cause off-target effects. We observed that after EGF treatment, EGFR was heavily phosphorylated (red) as expected, and EGFR co-receptors showed increased phosphorylation including ErbB2/Her2 (orange), ErbB3/HER3 (purple), MerTK (green), and EphB2 (yellow) (36–38). Interestingly, while EGFR phosphorylation was not sensitive to the presence of the peptide, phosphorylation of MerTK and EphB2 appeared to show a decrease with EGFR+PET1 treatment. EGFR-mediated phosphorylation of co-receptors may be one reason for changes in downstream signaling in the latter cases.

### PET1 separates the EGFR TM dimer by binding both helices simultaneously

Once we determined that PET1’s effect on cell migration was due to binding between PET1 and EGFR, we sought to further understand how PET1 binds the EGFR TM through molecular dynamics simulations (Figure 5). We employed Coarse Grain (CG) molecular simulations for our study to observe the binding of PET1 with the EGFR TMs. For this, we studied the association of EGFR-TM regions in the absence and presence of PET1 (Figure 5A, B). As the starting configurations, we modeled the membrane embedded regions as ideal monomeric helices placed in random orientation 5 nm apart from each other in a POPC membrane bilayer. The CG simulation was run for 4 µs in quadruplicates to assess the consistency of the results. In the case of the EGFR TM alone (Figure 5C), we saw that the two helices came together within 0.5 µs. Table S1 shows the PREDDIMER analysis for the EGFR TM dimers in comparison with the NMR structure (PDB ID: 5LV6) (5). In presence of PET1, the Fscor –a score reporting on the close and energetically strong packing of helices-were very low for the most populated cluster centers (two with 0, the two lesser populated structures with 3.5 and 2.6 respectively; Table S1) suggesting weak association between the EGFR TMs for most of the simulations. However, in the control simulations without PET1 the Fscor were high (3.2 to 4.0) suggesting stable interaction between the EGFR TMs. In the absence of PET1, the EGFR TMs assume a configuration which on average resembles that of the NMR structure thought to be the active state (RMSD of top cluster 1.7 Å, Table S1). This result, as well as CG simulations for other single membrane crossing TM receptors gives us confidence that the employed Martini 3.0 potential function leads to accurate EGFR TM dimer structures.

**Figure 5:**
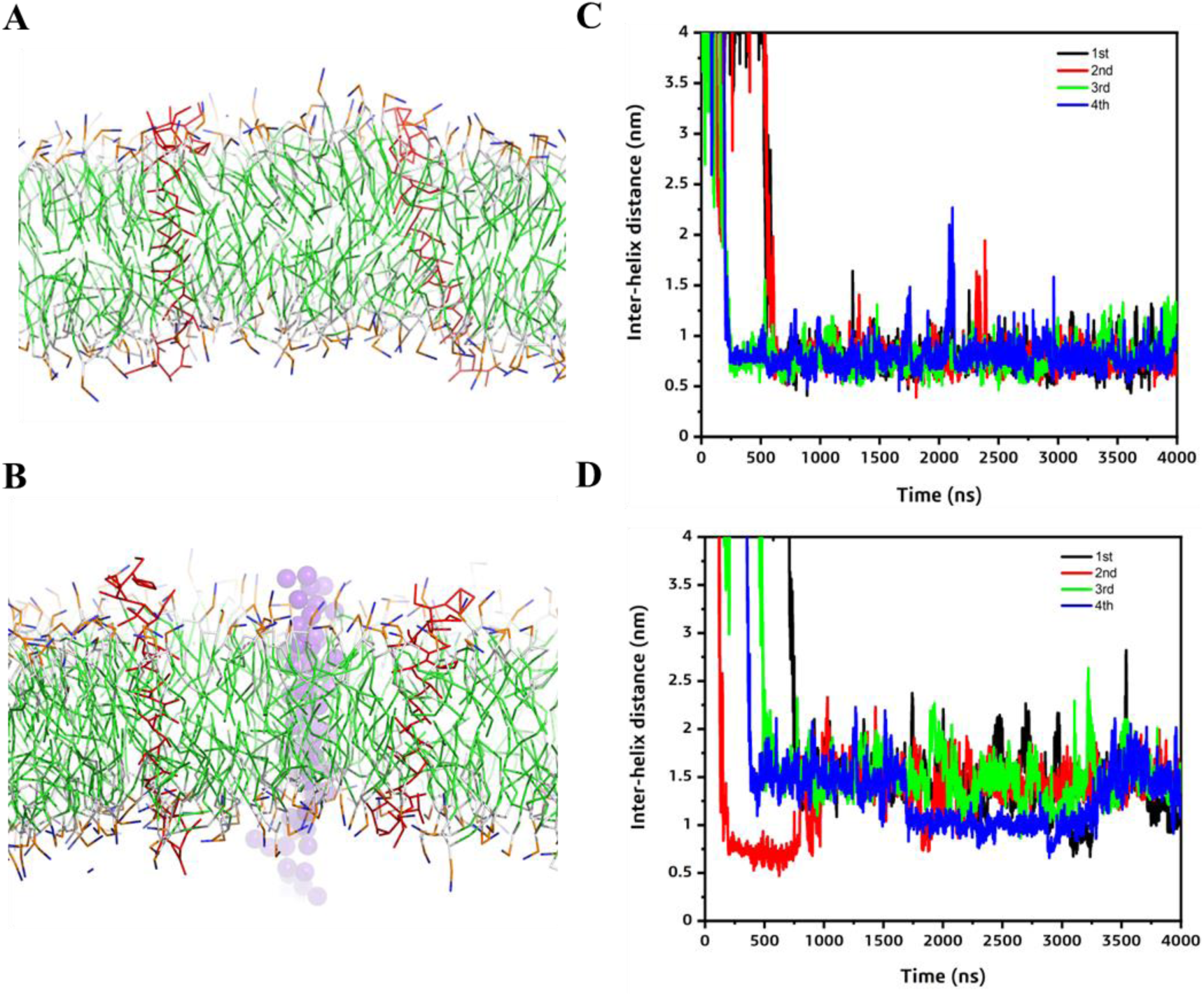
Molecular dynamics simulations. *Left panel*-Initial set up for the EGFR TM-only (**A**) and the EGFR-PET1 (**B**) systems in the lipid bilayer (solvent and ions not shown for clarity). The EGFR TMs and the PET1 are shown as red lines and purple spheres, respectively. *Right panel*-Inter-helical distance (COM) plots showing the association between the TM regions of EGFR in absence (**C**) and in the presence of PET1 (**D**). 1^st^, 2^nd^, 3^rd^ and 4^th^ simulation results are shown black, red, green and blue lines, respectively.

For the simulations with PET1, we assumed the simplest possible scenario, where PET1 binds to the EGFR TM dimer with a 1:2 stoichiometry. We observed the association of the PET1 monomer with the EGFR TM helices forming a heterotrimer within 1 µs in the four molecular dynamics trajectories (Figure 5D). To better observe the effect of PET1 association, we RMSD-clustered the simulations and superimposed the main conformer in Figures 6A, B. We also show the contact maps in Figures 6C, D, as averages over all simulations and plotted as the inter-helical distance between the EGFR TMs (considering the center of mass, COM, of the TM region) vs. the crossing angle between the two EGFR TM helices in Figs. 6E, F, yielding a 2D population map. In all cases the helices aligned close to parallel. In the EGFR TM dimer structure, the center of mass (COM) of the helices was spaced 0.65-0.95 nm, while with PET1 the distance increased to 0.90-1.15 Å or 1.30-1.55 nm. Thus, overall the presence of PET1 prevents the direct association of the EGFR TM helices for most of the time in the simulations as shown in Figure 6B. This is true in 3 of the 4 cluster centers but in the case of the third cluster center PET1 has a tilted orientation in the bilayer and associates more on the side of a weak EGFR TM dimer (Figure 6B), leading to the higher Fscor, as mentioned above. We also plotted the configurational transition of EGFR TM dimers in the absence and presence of PET1 (Supp. Figure 5), demonstrating that both systems are somewhat dynamic, especially the EGFR TM configurations in presence of PET1 which undergoes considerable fluctuations, most of which are accompanied by temporary increases in helix-to-helix COM distances.

**Figure 6.**
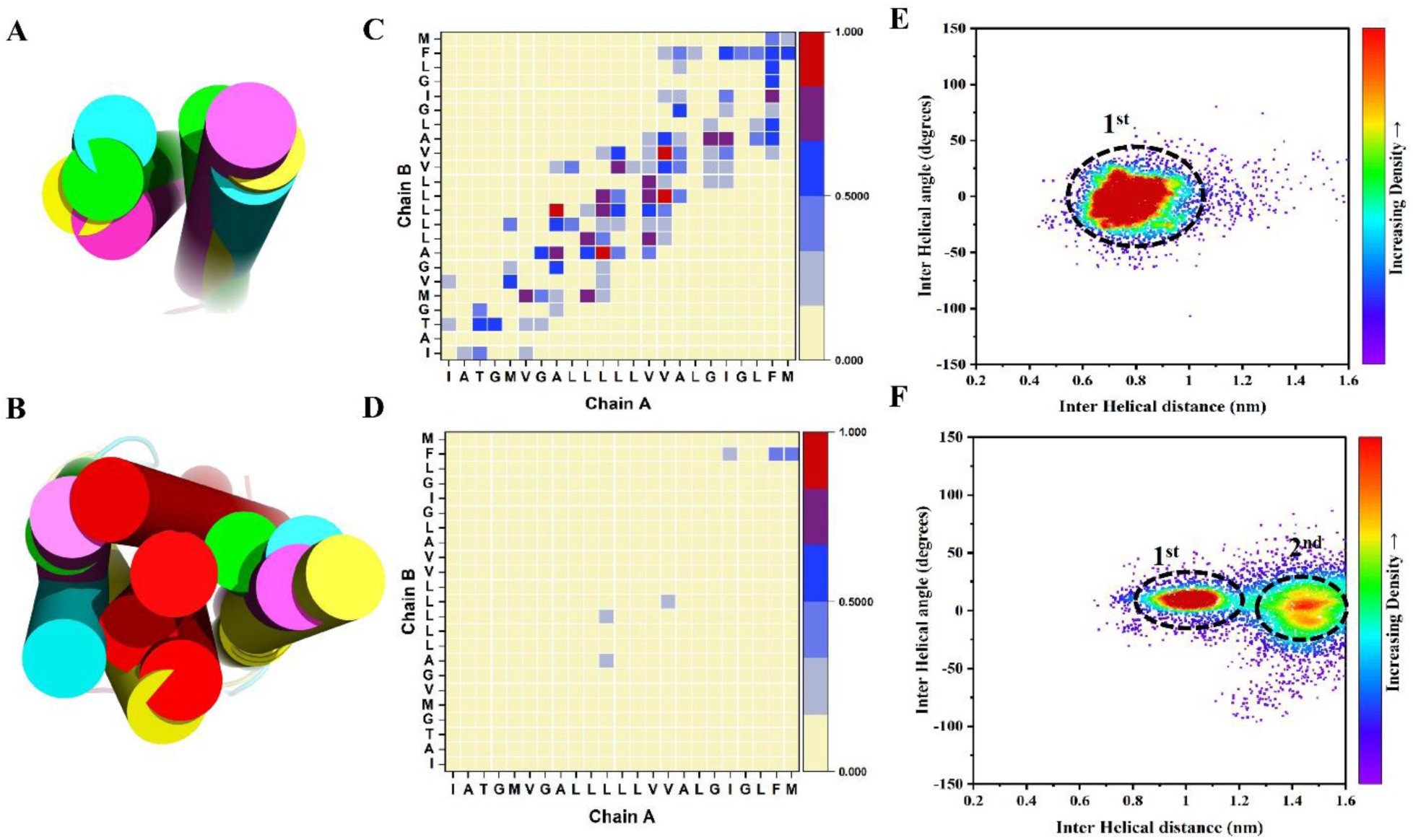
Comparison of the association of the EGFR TMs regions in the absence and presence of PET1. *Left panel*-Superimposition of the central conformers for all the four simulations in case of EGFR TM-only (**A**) and EGFR-PET1 (**B**) systems. PET1 is shown as red, and the EGFR TMs are in different colors. *Middle panel*-Simulation average contact map interface between the EGFR TMs for EGFR TM-only (**C**) and EGFR-PET1 (**D**) systems. Data from the last 1 µs simulations are considered for all the 4 simulations. Contact maps are calculated with a cut off of 5 Å. The color scale (yellow to blue to red) indicates the fractional occupation of TM contacts (0 to 1). *Right panel*-2D distribution plot (interhelix angle vs. distance) between the EGFR TMs for EGFR TM-only (**E**) and EGFR-PET1 (**F**) systems.Distance range clusters are indicated. Data from the last 1 µs simulations are considered for all the 4 simulations Corresponding data for a scrambled version of PET1 are shown in Fig S6.

As a control, we repeated the CG simulations with the scrambled PET1 peptide (SP) (Supp. Figure 4). The CG results showed that the SP was able to bind to the EGFR TMs (Supp. Figure 6), but this interaction did not robustly separate the EGFR dimer. In the presence of SP, the TM helices largely remained within ∼0.75 nm (Supp. Figure 6C), similarly to the case when the simulations were run in the absence of peptide (Figure 6E). Additionally, SP did not prevent key residues in the EGFR TMs to engage in dimer-stabilizing interactions (Supp. Figure 6B). These simulations suggest that the effect of PET1 is specific.

To benchmark the CG simulations, we applied AlphaFold-Multimer (39) to the complex formed by the EGFR TM helices and PET1 in a 2:1 stoichiometry. The artificial intelligence program predicted that PET1 binds to both TMs and blocks their C_t_ association (Supp. Figure 7), in agreement with the CG simulations. As a control, we also applied AlphaFold-Multimer to the isolated EGFR TM helices. The obtained prediction is in strong agreement with the TM structure believed to correspond to the active conformation of the receptor, solved by NMR (Supp. Figure 7) (40). The confidence score of both predictions had reasonable values (0.58 vs 0.43). However, the AlphaFold-Multimer prediction with SP yielded a low score (0.3) and is therefore not considered a robust prediction. To summarize, AlphaFold-Multimer and the CG simulations support the notion that PET1 disrupts native interactions between the TM helices of EGFR.

**Figure 7:**
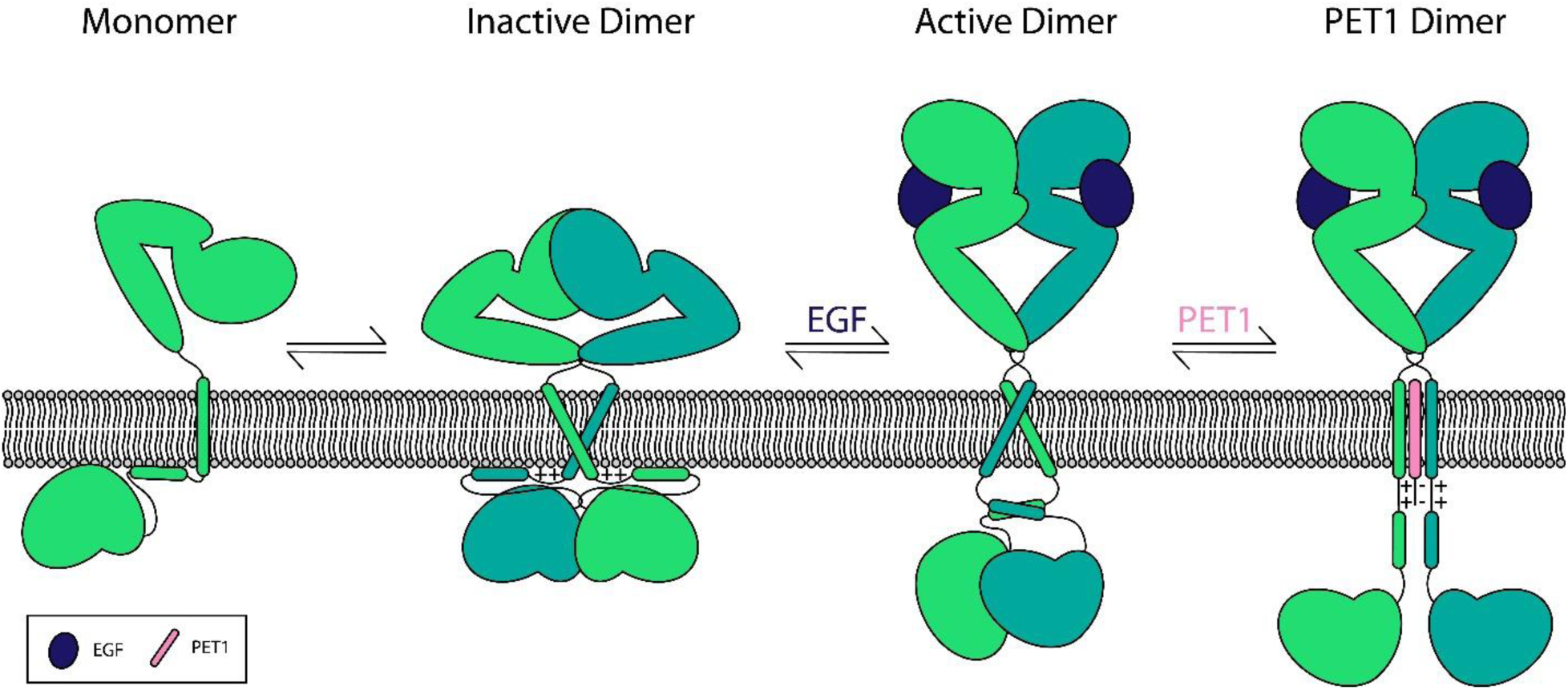
Model of the EGFR configurational changes caused by EGF and PET1. It is well established that EGFR exists in a monomer dimer equilibrium in its inactive state. The addition of EGF causes a configurational rearrangement for the extracellular, TM, JM, and kinase domains that allow autophosphorylation and activation. We propose that PET1 induces a configuration of the protein in which the TM interaction modifies the ability of the JM and kinase domain to arrange correctly for signaling to occur.

## Discussion

Our data indicate that PET1 is a pH-responsive peptide that inhibits EGFR activation through disruption of the EGFR TM dimer. The pH-responsive nature of PET1 is an important aspect of its design. Acidity-responsive peptides like pHLIP can preferentially target tumors over healthy tissue in mouse models, due to the more acidic environment of cancer cells (18–22). While EGFR is commonly overexpressed in cancer, there is still expression in healthy tissue. Thus, designing PET1 as a pH-responsive peptide will likely ameliorate any eventual off-target effects. We observed that PET1 displayed an increased pH_50_ in the presence of TM-EGFR (Figure 1D). However, the reported pH_50_ of PET1 in the presence of TM-EGFR is lower than the extracellular pH of tumors (pH 6.4-6.8). One must consider that pH_50_ determination by an N_t_ NBD tag consistently provides a lower pH_50_ value than determination by tryptophan spectral max, center of mass, or CD (24). Using this difference of 0.5, we can correct the determined pH_50_ in the presence of TM-EGFR to what is likely a more accurate pH_50_ of 5.85, which is closer to that of cancer cells. In addition, the pH_50_ was determined in pure POPC lipid vesicles. It is known that the lipid composition of a membrane can strongly affect the pH_50_ of pH-responsive peptides (18). Therefore this *in vitro* model membrane may not directly recapitulate the insertion conditions of the peptide into mammalian cell membranes (41). Regardless, our cellular results show that PET1 is able to insert into the plasma membrane and interact with EGFR at physiological pH, similarly to results obtained for the TYPE7 for the modulation of the EphA2 receptor (15).

Our model of PET1 inhibition of EGFR is shown in Figure 7. In its ligand-free form, EGFR primarily exists in an equilibrium between a monomer and dimer. Ligand binding shifts the equilibrium towards the dimer and induces a configurational change that allows kinase activation. In the inactive dimeric configuration, NMR reveals that the EGFR TM dimer has a helix-helix crossing angle of 30° utilizing the C_t_ AxxxG motif (PDB 2M0B) (5). This configuration causes the C_t_ ends of the TM to be positioned only 7.2 Å apart (4, 42, 43). The JM is therefore held too close together to form the antiparallel dimer characteristic of the active form, and the positively charged residues of the JM interact with the negatively charged lipids of the inner membrane leaflet (26, 41, 43). These electrostatic interactions hold the kinase domain close to the membrane where it is inactive (26). In contrast, binding of a ligand such as EGF causes the TM dimer to shift positions such that it dimerizes through the SxxxGxxxA motif at the N_t_ of the TM domain, with a helical crossing angle of −42° (4, 5, 40). Moreover, the N_t_ motif is on the opposite side of the helices, compared to the C_t_ AxxxG motif-which is utilized in the inactive state, thus leading to a rotation of both helices by 180° for receptor activation (Figure 7). In the active dimer the C_t_ end of the helices are 20 Å apart, a sufficient distance to allow JM antiparallel dimerization, which promotes asymmetric dimerization and activation of the kinase domain (41, 42). Our MD data in the presence of PET1 reveals a TM configuration unlike either the active or inactive state; PET1 is sandwiched between the two EGFR helices, which disallows most intramolecular contacts between the EGFR TM regions, and forces the center of mass of the TM regions to be approximately 15 Å apart. Furthermore, in this configuration, the negatively charged amino acids just outside the membrane on the PET1 sequence likely interact with the positive residues in the JM region of EGFR (not examined here by modeling, since only the very N_t_ region of the JM was included in the simulations). This is supported by our tryptophan fluorescence data (Figure 1E), which shows the EGFR JM residues in a more solvent exposed position in the presence of PET1. It is possible that PET1 forces the JM away from the membrane, due to an electrostatic attraction between the acidic residues of PET1 and the basic residues of the JM. This interaction might force a conformation that is neither close enough to force the JMs apart and to bind with the membrane, nor far enough to allow the JM to dimerize. We therefore propose that PET1 promotes a conformation that is different to the ligand bound or unbound dimer, and therefore disallows the native downstream effects of EGFR.

Indeed, the model proposed here is supported by previous findings. Prior MD simulations revealed that the EGFR TM is likely able to exist in configurations other than the active or inactive ones discussed above (44, 45). Additionally, cryo-EM has shown that another EGFR ligand, TGF-α, induces an extracellular conformation that is different than EGF’s and likely causes an intermediate TM conformation somewhere between the two previously discussed (46). This is further supported by crosslinking experiments in live cells that reveal that the EGFR TM-JM region is configured differently by binding of each of the 7 known ligands (45). Together, these experiments suggest that the TM dimer is more dynamic than originally thought, and it is further proposed that these small dynamic changes in the TM dimer exert large effects on the intracellular configurations and signaling of EGFR (45). Therefore, it is not at all unreasonable to expect that disruption of TM dimerization has a strong effect on EGFRs activity.

Intriguingly, PET1’s mechanism of activity is opposite that of TYPE7, the only other pH-responsive peptide published to date that targets an RTK (EphA2) (15). TYPE7 works by “stapling together” both TM helices of the EphA2 dimer, which stabilizes the ligand bound conformation and promotes downstream EphA2 signals (15). In contrast, PET1 disrupts TM binding to inhibit downstream EGFR signals. The opposing mechanism is interesting, as the peptides were similarly designed, with the E mutations placed on the helix interface that participates in ligand-independent dimerization. In contrast, the EphA2 TM homologous peptide N3, a variant of TYPE7 with the E residues placed on the interface that participates in ligand-dependent signaling, appears to function more similarly to PET1 (23). N3 also sits between the two helices and disrupts the EphA2 dimer entirely. As discussed above, it appears that disruption of the dimer only affects the ability of the RTK to be ligand-activated, not the basal levels of activity. Further work will be needed to understand how to fine-tune the design, so as the peptide stabilizes or disrupts a specific dimer conformation.

Our work is a proof-of-principle study that shows that targeting the TM of EGFR can lead to an efficient inhibition of this receptor. We describe the molecular mechanisms through which PET1 functions, in which PET1 biases the dimerization of the EGFR TM domain to allosterically regulate downstream function. Future work would benefit from fully characterizing the cellular mechanism of PET1: what phosphorylation patterns and signaling pathways are affected? Are other cell phenotypes besides migration affected? In addition, optimization of the peptide’s ideal concentration, half-life, and kinetics would be invaluable for better understanding PET1’s function. With further exploration of PET1, we might find that we have a new way to therapeutically target EGFR-mediated cancers that combats off-target effects and drug resistance.

## Methods

### Reagents and Peptides

Peptides (PET1, TM-EGFR, pHLIP, and Scrambled) were synthesized by Thermo Fisher Scientific (Waltham, MA) at ≥95% purity. Peptide purity was assessed by matrix-assisted laser desorption ionization-time-of-flight (MALDI-TOF) mass spectrometry. The matrix α-cyano-4-hydroxycinnamic acid (α-HCCA) and trifluoroacetic acid (TFA) were purchased from Sigma-Aldrich (St. Louis, MO). Sodium phosphate and sodium acetate buffers were also purchased from Sigma-Aldrich (St. Louis, MO). Succinimidyl 6-(N-(7-nitrobenz-2-oxa-1,3-diazol-4-yl)amino) hexanoate (NBD-X,SE) and DyLight 680 NHS-ester were purchased from Thermo-Fisher Scientific (Waltham, MA). Anti-EGFR (D38B1) XP® Rabbit mAb #4267 and anti-EGFR Mouse mAb (IP Specific) #2256 were purchased from Cell Signaling Technology (Danvers, MA). The anti-β-actin antibody was purchased from Abcam (Cambridge, MA). Secondary IRDye® (680RD and 800CW) Goat anti-Rabbit and anti-Mouse were purchased from LI-CORE (Lincoln, Nebraska). Secondary Alexa Fluor 488 anti-Rabbit dye was purchased from Thermo-Fisher Scientific (Waltham, MA).

### Peptide Dye Conjugation

For NBD and DL680 conjugation of PET1, the esterified version of each dye (Thermo Fisher) was linked to the N-terminus. Dye suspended in DMF was added to PET1 dissolved in 100 mM sodium phosphate, 150 mM sodium chloride (pH 7.0) at a dye to peptide molar ratio of approximately 1:10 for NBD and 1:5 for DL680. The mixture was shaken for 1.5 hr and then centrifuged at 14,000 x g to remove precipitated dye. The supernatant was then run on a PD10 desalting column with 1 mM sodium phosphate (pH 7.2) buffer to separate free dye from conjugated peptide.

### Matrix-assisted laser desorption/ionization – time of flight (MALDI-TOF)

Conjugation efficiency was determined using MALDI-TOF. Peptides were added to a saturated solution of α-cyano-4-hydroxycinnamic acid in 70% methanol, 0.05% TFA and dried onto the MSP AnchorChip target plate (Bruker, Billerica, MA) using the dried droplet method. The Bruker Microflex MALDI-TOF mass spectrometer was calibrated with the Bruker Peptide Calibration Standard II (Billerica, MA). Mass spectra were analyzed using FlexAnalysis software (Bruker, Billerica, MA).

### Liposome Preparation

Lipids were purchased from Avanti Polar Lipids, Alabaster, AL. 1-palmitoyl-2-oleoyl-sn-glycero-3-phosphocholine (POPC) stocks were suspended in chloroform. Aliquots were dried under a stream of argon gas and then subjected to vacuum at least two hours before resuspension in 10 mM sodium phosphate (pH 7.4). For proteo-liposomes containing TM-EGFR, stocks of TM-EGFR in methanol were mixed with POPC prior to drying. Drying was performed in 13 mm glass culture tubes that had been piranha (75% H_2_SO_4_, 25% H2O2) cleaned for three minutes to reduce peptide sticking to the glass. Resuspended samples were extruded using a Mini-Extruder (Avanti Polar Lipids, Alabaster, AL) through a 100 nm membrane (Whatman, United Kingdom) to form large unilamellar vesicles (LUVs).

### Circular Dichroism (CD)

Stocks of POPC and PET1 were prepared in chloroform and 1 mM sodium phosphate (pH 7.4) buffer, respectively. An aliquot of POPC was dried under a stream of argon gas before placed in a desiccator for at least 2 hours. The POPC film was resuspended with 1 mL of 1 mM NaPi pH 7.4 buffer and extruded through a 100 nm Nuclepore Track-Etch Membrane (Whatman, United Kingdom) to produce large unilamellar vesicles (LUVs). PET1 was diluted to a working concentration of 7 μM peptide suspended in 20 mM sodium phosphate pH 7.5 or 20 mM sodium acetate pH 4.3. PET1 was incubated with LUVs at a 150:1 lipid to peptide molar ratio. Samples were recorded on a Jasco J-815 CD spectrometer using a 2 mm quartz cuvette (Starna Cells Inc., Atascadero, California). All conditions were averaged over 2 technical replicates. Appropriate buffer backgrounds were collected on the same day and subtracted appropriately.

### Oriented Circular Dichroism (OCD)

Stocks of POPC and PET1 were suspended in chloroform and 1,1,1,3,3,3-Hexafluoro-2-propanol (HFIP), respectively. An aliquot of POPC was dried under a stream of argon gas before placed in a desiccator for at least 2 hours. The POPC film was resuspended with a calculated volume of PET1 stock solution to reach a 50:1 lipid to peptide molar ratio and dried correspondingly. The POPC-PET1 film was resuspended with HFIP and deposited homogenously across two circular quartz slides (Hellma Analytics, Germany) cleaned with piranha solution. These slides were placed in glass petri dishes and balanced horizontally within a chemical hood at room temperature overnight to ensure complete HFIP evaporation. Lipid films on each slide were rehydrated with 150 mL of 100 mM sodium acetate buffer pH 4.24 for 16 hours in a 96% relative humidity chamber packed with saturated K2SO4. The majority of buffer was removed, and the slides were assembled into an OCD cell packed with saturated K2SO4 to maintain humidity. The OCD spectra were recorded on a Jasco J-815 CD spectrometer and averaged over eight 45° rotations of the cell. POPC lipid backgrounds were collected separately and subtracted appropriately.

### pH Titration Assay

LUVs and proteoliposomes prepared as above in 10 mM sodium phosphate (pH 7.4), were incubated for at least an hour with PET1-NBD at a lipid:TM-EGFR:PET1 molar ratio of 1000:5:1. Stocks were then diluted into a series of 100 mM sodium phosphate or sodium acetate buffers at pH’s between 4 and 7.6 in 0.2 intervals. The NBD fluorescence spectra were recorded at 25°C with excitation at 470 nm and an emission range of 520 – 600 nm using a Cytation five imaging plate reader (Biotek Instruments, Winooski, VT). Lipid blanks were prepared at the highest and lowest pH, averaged, and subtracted from the test spectra. The fluorescence (I_i_) and wavelength (λ) of each curve was used to calculate the center of mass (CM) at each pH using Equation 1.

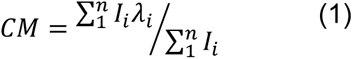

The CM was plotted against pH to determine the pH_50_ using Equation 2.

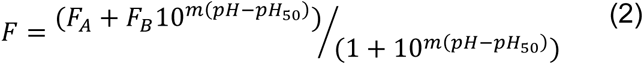

### Tryptophan Fluorescence Assay

LUVs and proteoliposomes were prepared as above. Appropriately pH adjusted 100 mM sodium phosphate (pH 7.4) or sodium acetate (pH 4.3) buffer and PET1 were added to LUVs for a final concentration of 200 μM POPC, 1 μM TM-EGFR, and 5 μM PET1. Samples were incubated for 1 hr at room temperature (19-21℃) to allow peptide binding to come to equilibrium. Tryptophan fluorescence spectra was then obtained on a Cary Eclipse Fluorescence Spectrophotometer at an excitation wavelength of 280nm (Agilent Scientific, Santa Clara, CA). For all treatments lipid blanks were subtracted.

### Cell Culture

A375, A431, and Cos7 cell lines were obtained from ATCC (Manassas, VA) and maintained at 5% CO_2_ and 37°C in Dulbecco’s Modified Eagle’s Medium (DMEM) supplemented with glucose, 10% fetal bovine serum (FBS), and 100 U/mL penicillin-streptomycin. Cells were passed at 80% confluency and were not used beyond 40 passes. Cell lines were STR tested for authentication via ATCC.

### Co-localization

A431 cells were plated at ∼80% confluency on a #1.5 glass coverslip, allowed to adhere for 24 hours, and then starved overnight. Cells were then treated with serum-free DMEM without (No Treatment) or with (PET1) PET1-DL680 for 1 hour prior to a 5 min EGF 100 ng/mL treatment. Cells were washed with PBS containing 1 mM MgCl_2_ and 100 mM CaCl_2_ (PBS++), fixed at 37°C for 15 minutes in 4% paraformaldehyde, and permeabilized for 10 minutes at room temperature with 1% Triton X-100. Cells were blocked with 3% BSA, and primary anti-EGFR XP (1:100) antibody was incubated overnight in 1% BSA at 4°C. Cells were washed, and secondary anti-rabbit conjugated to Alexa-fluor 488 (1:1000) was incubated 1 hr at room temperature. Cells were then stained for DAPI (1 µg/mL) for 5 minutes and mounted to a microscope slide using Diamond Anti-fade mounting media. After curing, cells were imaged using a 63x 1.4NA oil objective on a Leica SP8 White Light Laser Confocal Microscope. Images were the product of 3-fold line averaging. Three to five images were taken per coverslip and Pearson’s Correlation Coefficient (r) was calculated via the Coloc2 plugin from ImageJ.

### Co-immunoprecipitation

A431 cells were plated in a 12 well plate to ∼80% confluency and allowed to adhere for 24 hours. Cells were then starved overnight and treated with serum free-media without (No Treatment) or with (PET1) PET1-DL680 at 1, 2 and 5 µM for one hour before a 5 minute treatment with or without EGF (100 ng/mL). Cells were then washed, scraped from the plate using Co-IP buffer (50 mM tris-HCl (pH7.4), 150 mM NaCl, 5 mM EDTA, and 1% NP40) containing protease and phosphatase inhibitors, and allowed to sit on ice 30 minutes prior to a 10 min centrifugation at 13,000 x g. The pellet was discarded and 100 ng of total protein was diluted to 400 µL with anti-EGFR IP (1:100) antibody and rotated at 4°C overnight. 60 µL of pre-washed Protein A magnetic beads (Cell Signaling) were added and rotated for 2 hr. Then lysate was removed and beads were washed 4 x 10 min at room temperature with Co-IP buffer. Protein was eluted at 100°C for 5 minutes in 2X Laemmli sample buffer containing no dye, and eluate was run on a 4-20% SDS-PAGE gel. The gel was imaged for 680 nm fluorescence on an Odyssey CLx Imaging System (LI-CORE, Lincoln, Nebraska) before being transferred to 0.2 µm nitrocellulose, blocked with 5% milk, and blotted overnight with anti-EGFR XP antibody (1:1000). The membrane was washed and blotted with IRDye 800CW anti-rabbit secondary, and imaged for 680 and 800 nm fluorescence using the Odyssey as above.

### MTS Toxicity

A375 cells were plated in a clear, flat bottom 96-well plate to 80% confluency and allowed to adhere for 24 hours. Cells were then treated with phenol free DMEM containing 10% FBS alone (No Treatment) or containing EGF (100 ng/mL), PET1 (2 µM), pHLIP (2 µM), or the peptides in combination with EGF. Treatments were incubated 2 hours before addition of the MTS (3-(4,5-Dimethylthiazol-2-yl)-5-(3-carboxymethoxyphenyl)-2-(4-sulfophenyl)-2H-tetrazolium) reagent (Thermo-Fisher Scientific, Waltham, MA) and incubation was continued another 1.5 hours. Finally, absorbance at 490 nm was read using a Biotek Cytation V microplate reader with Gen5 software (Winooski, VT).

### PIE-FCCS

Pulsed interleaved excitation fluorescence cross correlation spectroscopy (PIE-FCCS) was used to study the effect of PET1 on the lateral oligomerization of EGFR. Expression vectors from previous studies were used to label EGFR at the C-terminus with EGFP and mCherry.(47) These vectors were expressed in COS7 cells purchased from Sigma Aldrich. COS7 cells were cultured in DMEM (Calsson Lab, Smithfield, UT) supplemented with 10% FBS (Sigma Aldrich) and maintained in a humidified incubator with 5% CO_2_ at 37 °C. To prepare for PIE-FCCS experiments, the cells were split, seeded on to a 35-mm MatTek plate (MatTek Corporation, Ashland, MA), and incubated until the confluency reached ∼70%. The plasmid constructs were transiently co-transfected to COS7 cells using Lipofectamine2000 (Invitrogen) approximately 24 hours before the data acquisition. Data were recorded on live cells before and after peptide or ligand addition as described previously.(47, 48) The two-color PIE-FCCS experiment and the auto/cross-correlation analysis allow us to evaluate the expression density, diffusion (reported as an effective diffusion coefficient, D_eff_), and the oligomerization state (reported as fraction correlated, *ƒ_c_*). The density of the receptors expressed was in the range of 100-2000 receptors/µm^2^. Lower D_eff_ and higher *ƒ_c_* values indicate formation of larger oligomers. To test for the effect of PET1, 2 µM or 2 µg/mL of PET1 were added to the well with 2 mL imaging media and incubated for 10 min before data acquisition. Data were acquired up to 60 min after ligand addition.

### Cell Migration

A375 cells were plated to 50% confluency on a 10 cm dish and allowed to adhere for 24 hr before overnight starvation in serum free DMEM. Cells were then trypsinized with 0.05% trypsin for the minimal amount of time required to remove from the plate, washed, and brought to a density of 2 x 10^5^ cells/mL in serum free DMEM. 100 µL of cells were plated on the top of a 6.5 mm transwell polycarbonate membrane insert with an 8 µm pore size (Corning 3422) while 600 µL of DMEM containing 10% FBS alone (No Treatment) or in combination with EGF (100 ng/mL), PET1 (2 µM) or both EGF and PET1 together was in the bottom of the insert. Cells were allowed to migrate 24 hr at 37°C and 5% CO_2_ before chambers were washed, cells remaining on top of the membrane were scraped off, and cells on the bottom of the membrane were fixed with methanol and stained using hematoxylin and eosin. The membrane was then cut from the chamber and mounted to a microscope slide for imaging using a 10X objective on a Biotek Cytation V microplate reader with Gen5 software (Winooski, VT).

### RTK Array

A431 cells were blotted using an R&D RTK array kit (ARY001B). Cells were plated at 80% confluency on a 6 well plate, allowed to adhere 24 hr, and then starved overnight using serum free DMEM either alone or containing PET1 (2 µM). Cells were then incubated for 5 minutes with PET1 alone, EGF alone (100 ng/mL), or PET1 and EGF in combination. Cells were then washed and scraped from the plate using 1X kit lysis buffer and agitated at 4C for 30 minutes. Lysates were centrifuged 10 minutes at 13,000 x g and the supernatant was quantified using a DC assay kit (Bio-Rad, Hercules, California). 200 µg of total protein were diluted into a total of 1.5 mL Array Buffer and rocked over the pre-blocked array membrane containing various total RTK antibodies overnight at 4°C. After washing, anti-phospho-tyrosine antibody conjugated to HRP (kit) was blotted for 2 hr at room temperature, washed, and the HRP was developed using Chemi Reagent Mix (kit). The blots were imaged using a LI-COR Odyssey CLx imager, with 10 sec to 3 min exposure times.

### Modeling of the Transmembrane (TM) peptides

The NMR structure of EGFR TM dimer (PDB ID: 5LV6)(5) was obtained from www.rcsb.org. The TM region and the membrane proximal N-terminal residues of EGFR from P^641^-I^673^ was extracted from the NMR structure and the remaining modified TM C-terminal residues from C^674^WN^676^ were modeled as an extended conformation of amino acids (ϕ, ψ= ±120°) in PyMOL (The PyMOL Molecular Graphics System, Version 2.4. Schrödinger, LLC). The PET1 peptide [T^638^NGPKIPS<u>IATGMVGA**E**LLLGVVA**E**GIGLFM</u>**EEEE**^672^] was modeled as transmembrane helix from I^646^-M^668^ based on EGFR TM NMR structure as the template using Modeller (49) and the remaining N-terminal residues (T^638^-S^645^) and C-terminal residues (E^669^-E^672^) were modeled as an extended conformation (ϕ, ψ= ±120°) in PyMOL. This PET1 peptide has 7 mutations as shown in Figure 1A. The scrambled peptide (SP) [F^638^KLAAVNG<u>GIGSTGGI**E**MVIPGMP**E**LTLALV</u>**EEEE**^672^] was also modeled as TM helix from G^646^-V^668^ in a similar manner with the terminal residues as extended conformation.

### Coarse-grain (CG) molecular dynamics simulation

To check the influence of the PET1/SP on the dimerization of EGFR TMs, we built an initial system configuration which contains the two EGFR TM monomers and the PET1/ SP peptide, where all three monomer TM units placed perpendicular to the membrane and 5 nm apart from each other. As a reference for the above systems, we also ran simulations with the same system set-up but omitting the PET1/SP peptide, i.e. the two EGFR TM peptides by themselves as the control. The atomistic (AT) modeled systems with EGFR TMs: PET1 (2:1); EGFR: SP (2:1) and EGFR TMs alone, each placed as above, were converted to coarse-grained (CG) representation using the *martinize2.py* workflow module of the MARTINI3 forcefield(50) (version 3.0.4.28) considering the secondary structure DSSP assignment. CG simulations were performed using Gromacs version 2016.5 (51). The setting up of the POPC bilayer was done using the insane.py script (52) (typically 684 lipid and 19,300 CG water molecules for 2:1 peptide systems; 324 lipids and 4,350 CG water molecules for control EGFR TM dimer systems) around the peptides. The pH of the system was 4.5, setting all the Glutamate residues in the peptides to be protonated. The systems were equilibrated for 500 ps. The electrostatic interactions were shifted to zero between 0 and 12 Å and the Lennard-Jones interactions were shifted to zero between 9 and 12 Å. The V-rescale thermostat was used with a reference temperature of 320K in combination with a Berendsen barostat at 1 bar reference pressure, with a coupling constant of 1.0 ps, a compressibility of 3.0 × 10^-4^ bar ^-1^. The integration time step was 20 fs and all the simulations were run in quadruplicate for 4 µs.

### Data Analysis

Interhelix distances between the center of masses of the TM regions were calculated and the clustering were performed with a cut off 6Å using the module of the Gromacs by combining the entire trajectories of all the four simulations. PREDDIMER webserver was used for analyzing the TM dimers cluster centers based on the Fscor, helix crossing angle and helix rotation angle.(53) For the latter, the residue L657, which is part of the regular TM helix, was chosen and angles were read of PREDDIMER energy maps. The contact maps for the TM regions between the helices were calculated with a cut off 4 Å for all the backbone and side-chain atoms. Sequence alignments were done using ClustalX (54). Data were plotted in GraphPad Prism (version 6 for Windows, GraphPad Software, La Jolla California USA, www.graphpad.com).

AlphaFold-Multimer (39) was used to predict models for EFGR TMs alone and in complex with the PET1 or SP peptide (2:1).

## Data Availability

Data will be provided upon request.

## Supporting Information

This article contains supporting information.

## Acknowledgements

This work was supported by NIH grant R35GM140846 (to FNB), a National Science Foundation grant CHE-1753060 (to AWS) and a grant from the National Eye Institute R01EY029169 and previous grants from NIGMS (R01GM073071 and R01GM092851) to the Buck lab. The simulations were run at the local computing resource in the core facility for Advanced Research Computing at Case Western Reserve University. We thank Ryan J Schuck, Charles M Russell, Alyssa E Ward, and Yujie Ye for their thoughtful comments on this manuscript.

## Supplementary Materials

**Supplementary Figure 1:**
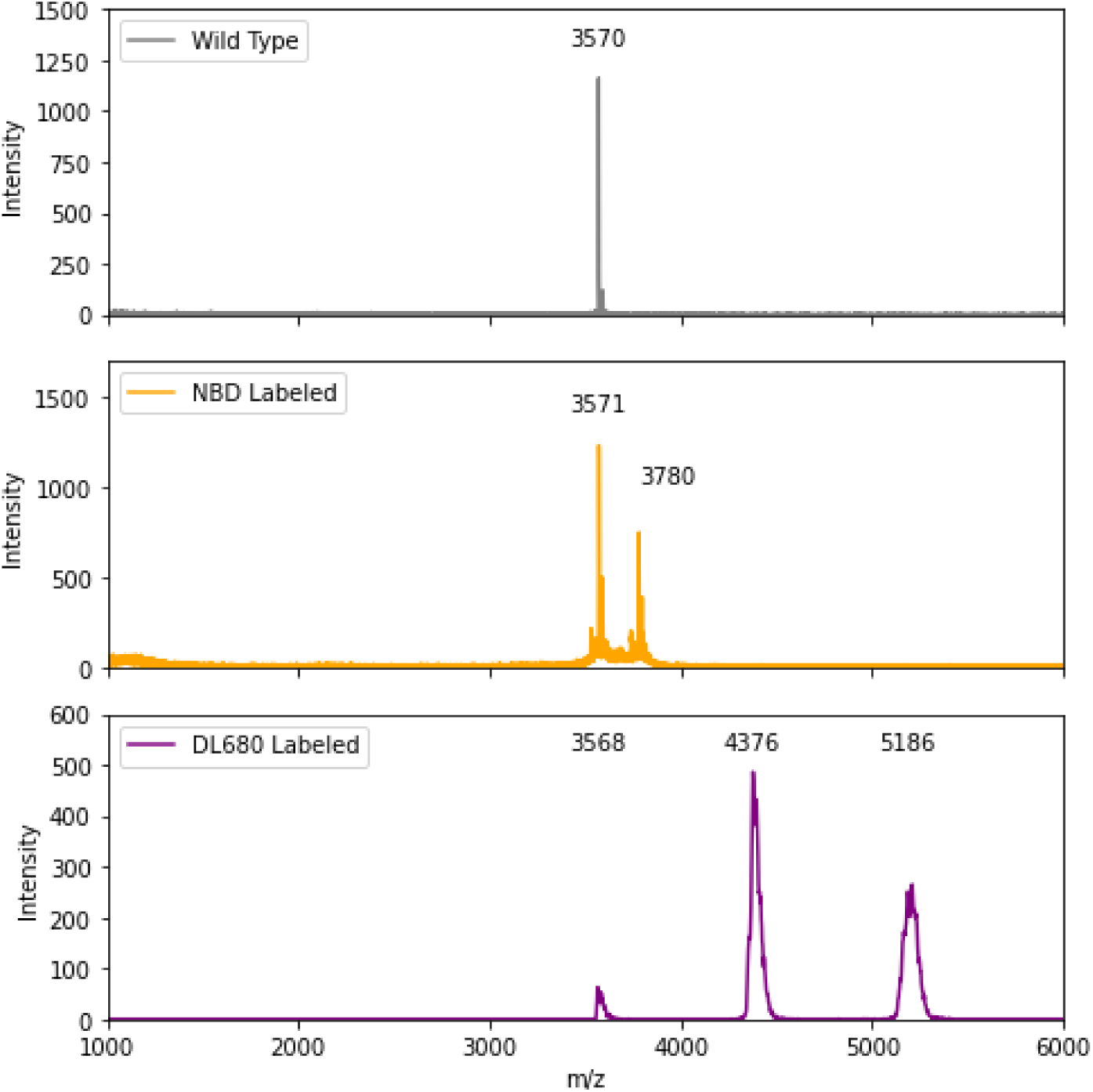
MALDI-TOF of PET1, PET1-NBD, and PET1 DL680.

**Supplemental Figure 2:**
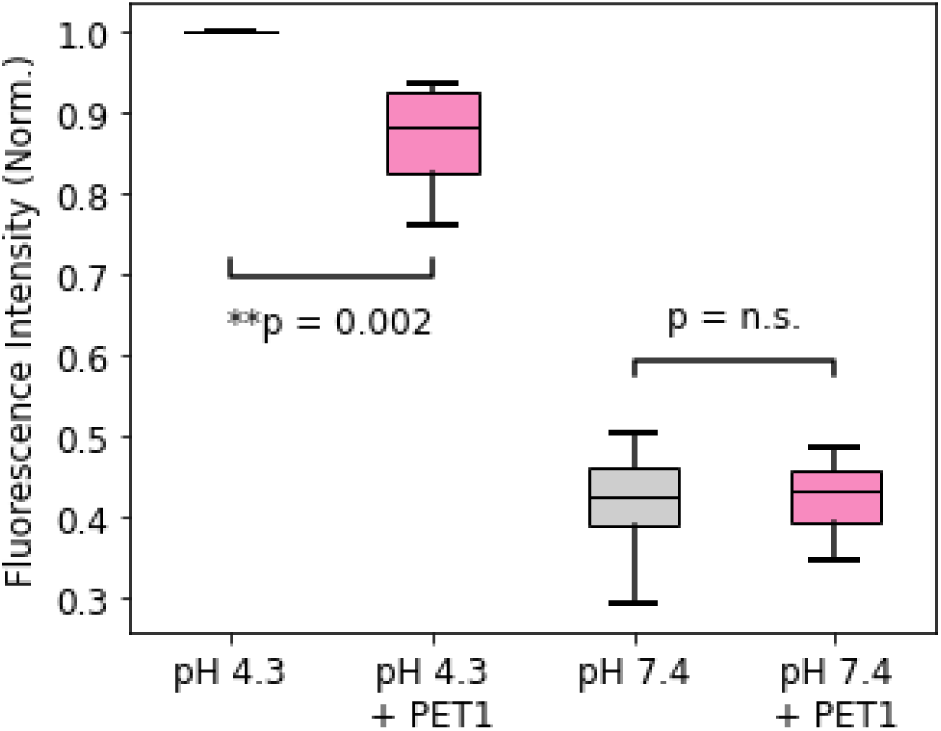
PET1 decreases the fluorescence intensity of TM-EGFR only at acidic pH. The fluorescence spectra of TM-EGFR in POPC lipid vesicles at pH 4.3 and 7.4 in the presence (pink) or absence (grey) of PET1. Box plot conveys the fluorescence at the max of the curve. N = 6 with each biological replicate normalized to pH 4.3 conditions. Statistical analysis was performed using a Kruskal Wallis test -H(3) = 19.75, *p* = 0.0002-with a Mann Whitney U test for comparisons between groups.

**Supplementary Figure 3:**
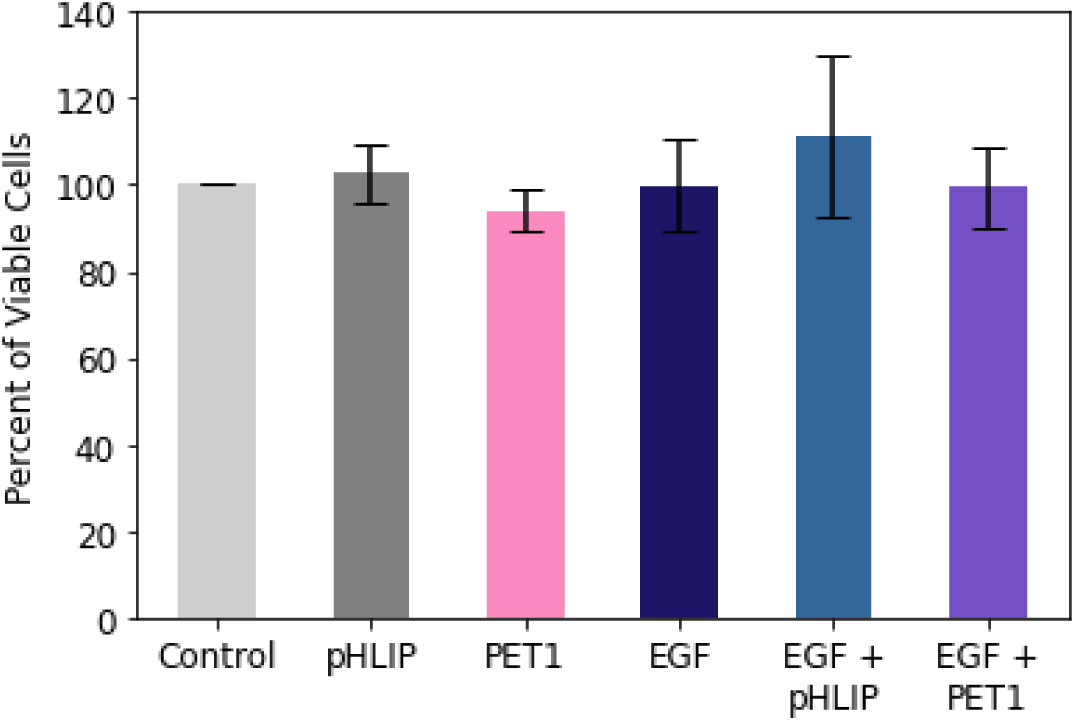
PET1 is not toxic to A375 cells. MTS assay was performed to evaluate cell viability. The pHLIP peptide was used as a control for a pH responsive peptide with no toxicity (16). Within each biological replicate the number of untreated cells was normalized to 100% viability. N = 3. Error bars denote standard deviation of the mean.

**Supplementary Figure 4:**
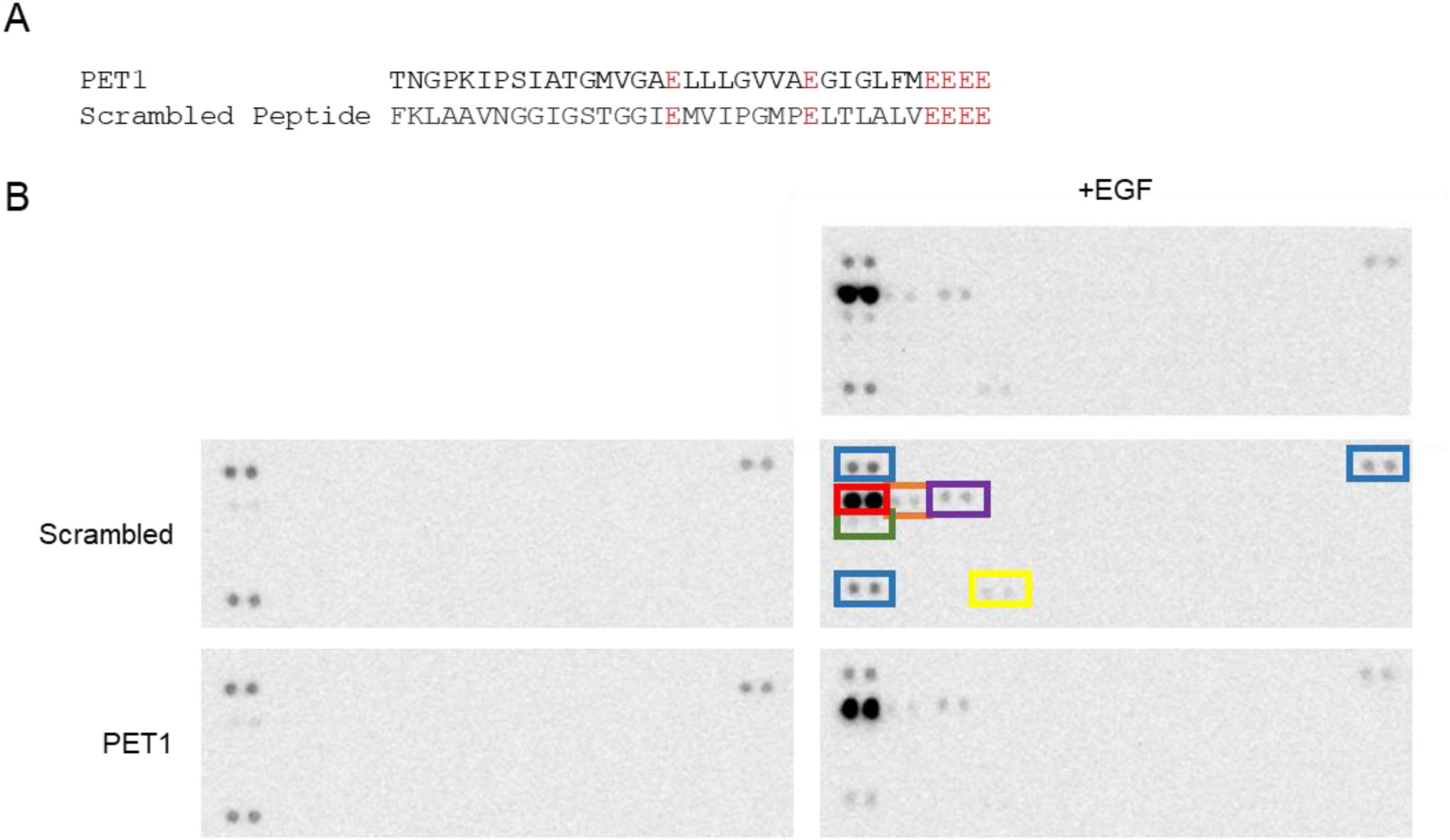
PET1 does not activate other RTKs. **A,** Sequence of PET1 peptide and scrambled peptide used as a control. E residues were kept constant and are highlighted in red. **B,** Lysates of A431 cells were treated with or without PET1 overnight followed by a 5 minute treatment with or without EGF. Tyrosine phosphorylation on a variety of RTKs was probed with a Human Phospho-RTK Array Kit. Three sets of reference dots (blue) allow determination of RTK identities. Specific RTK are boxed: EGFR (red), ErbB2 (orange), ErbB3 (purple), MerTK (green), and EphB2 (yellow). Other RTKs did not show phosphorylation in any conditions: ALK/CD246, Axl, DDR1, DDR2, Dtk, EphA1, EphA2, EphA3, EphA4, EphA5, EphA6, EphA7, EphA10, EphB1, EphB3, EphB4, ErbB2, ErbB3, ErbB4, ErbB6, FGFR1, FGFR2 alpha, FGFR3, FGFR4, Flt-3/Flk, HGF R/c-MET, IGF-I R, Insulin R/CD220, M-CSF R, Mer, MSP R/Ron, MuSK, PDGF R alpha, Ret, ROR1, ROR2, Ryk, PDGFRβ, SCF R/c-kit, Tie-1, Tie-2, TrkA, TrkB, TrkC, VEGF R1/Flt-1, VEGF R2/KDR, and VEGF R3/Flt-4. Blot identities can be found at https://resources.rndsystems.com/pdfs/datasheets/ary001b.pdf?v=20220613&_ga=2.236641154.205998430.1655143431-478466794.1651682163

**Supplementary Figure 5.**
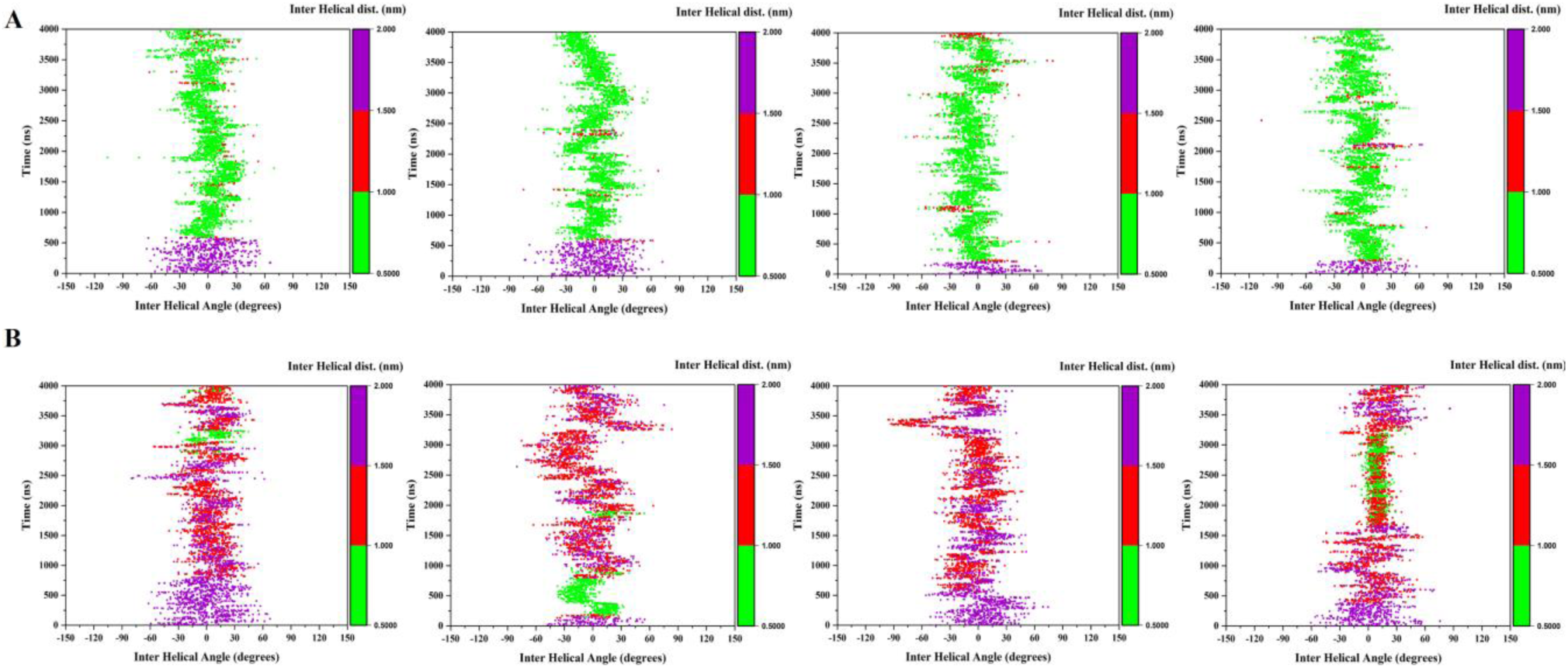
2D Plots showing the configurational transition of EGFR TM dimers over the simulation time considering the inter-helical angle vs inter-helical distance for the EGFR TM-only (**A**) and EGFR-PET1 (**B**). Results from all 4 trajectories are shown. The plots are colored based on the inter-helical distance from 0.5-1.0 nm (green), 1-1.5 nm (red) and 1.5-2.0 nm (purple).

**Supplementary Figure 6.**
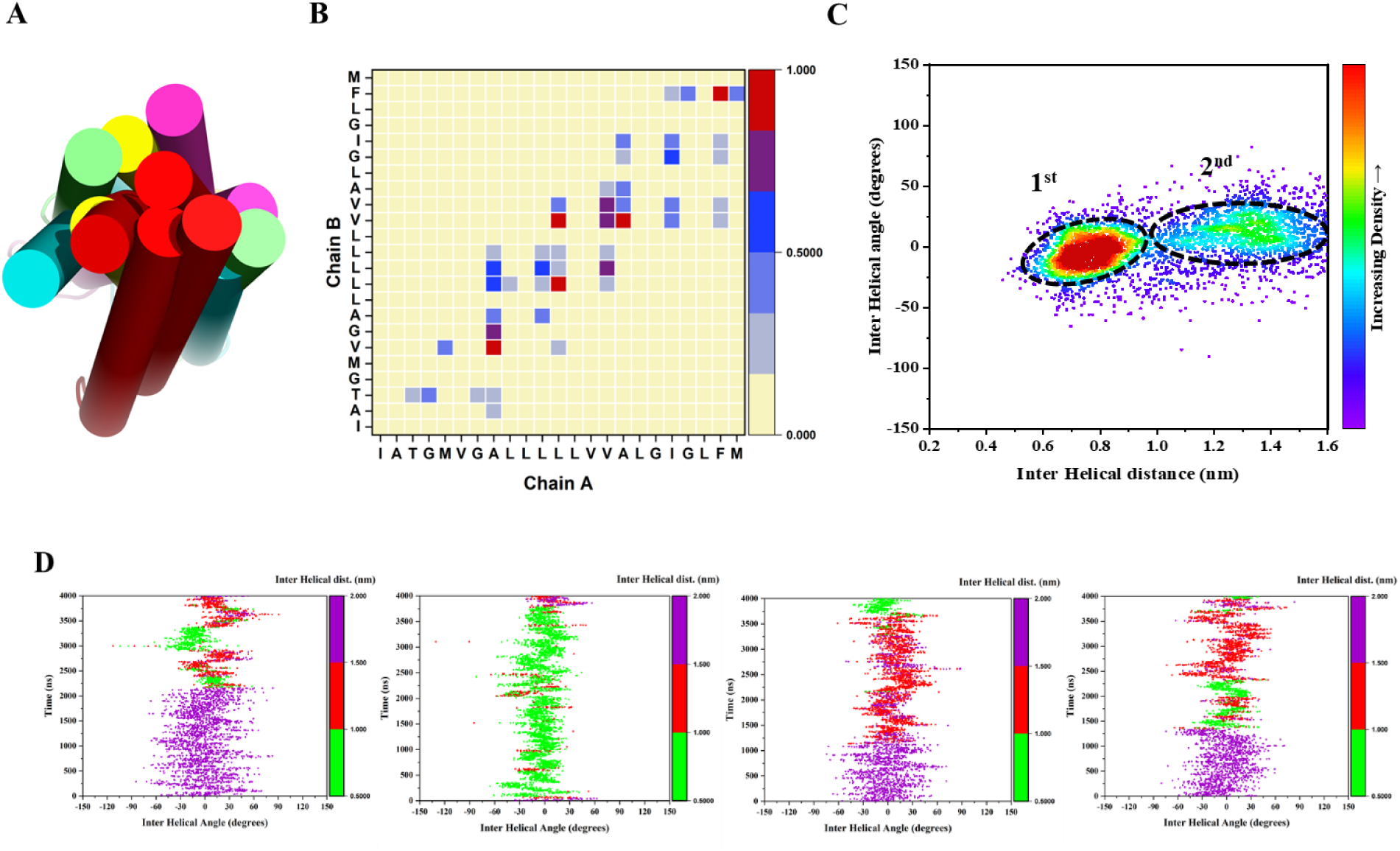
Association of the EGFR TMs regions in the presence of the Scrambled peptide (SP). **A,** Superimposition of the central conformers for all the four CG simulations in case of EGFR-SP systems. The SP peptide is shown in red and the EGFR TMs in different colors. **B,** Contact map interface between the EGFR TMs for EGFR-SP systems. Data from the last 1 µs simulations are considered for all the 4 simulations. Contact maps are calculated with a cut off 5 Å. The color scale (white to blue to red) indicates the fractional occupation of TM contacts (0 to 1). **C,** 2D distribution plot (interhelix angle vs. distance) between the EGFR TMs for EGFR-SP systems. Populated clusters are named as 1^st^, 2^nd^ and 3^rd^. Data from the last 1 µs simulations are considered for all the 4 simulations. **D,** Plots showing the configurational transition of EGFR TM dimers over the simulation time considering the inter-helical angle vs inter-helical distance.

**Supplementary Figure 7.**
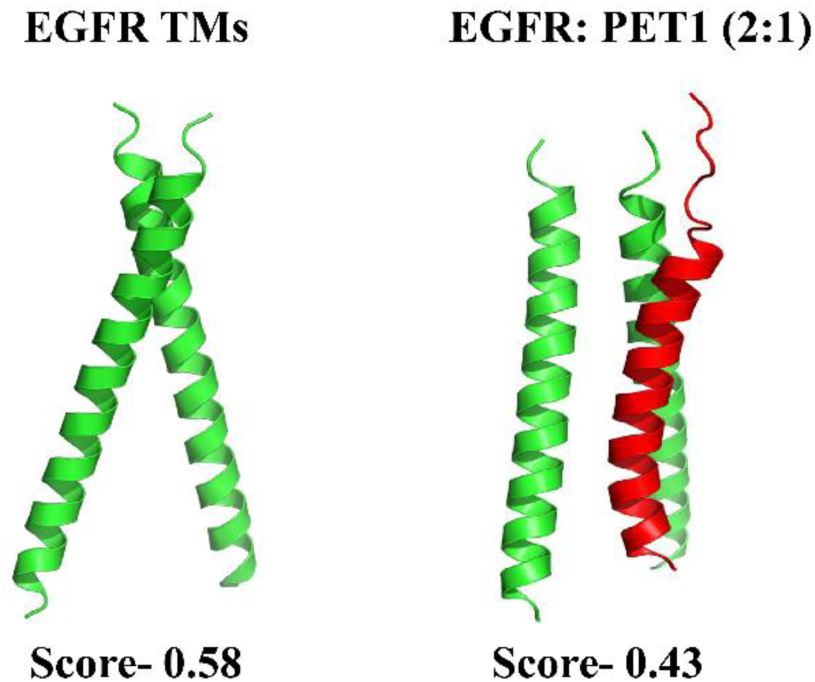
Comparison of AlphaFold-Multimer predicted structures of EGFR TM (green) homodimer and in complex with PET1 (red). We also predicted the structure of the EGFR: SP (2:1), but the AlphaFold-Multimer score was low (0.3). Since the native sequence has been randomly arranged in the SP, there are few homologous sequences available in the training dataset. Therefore, the SP prediction was poor and accuracy of that prediction is uncertain (data not shown).

**Table S1:**
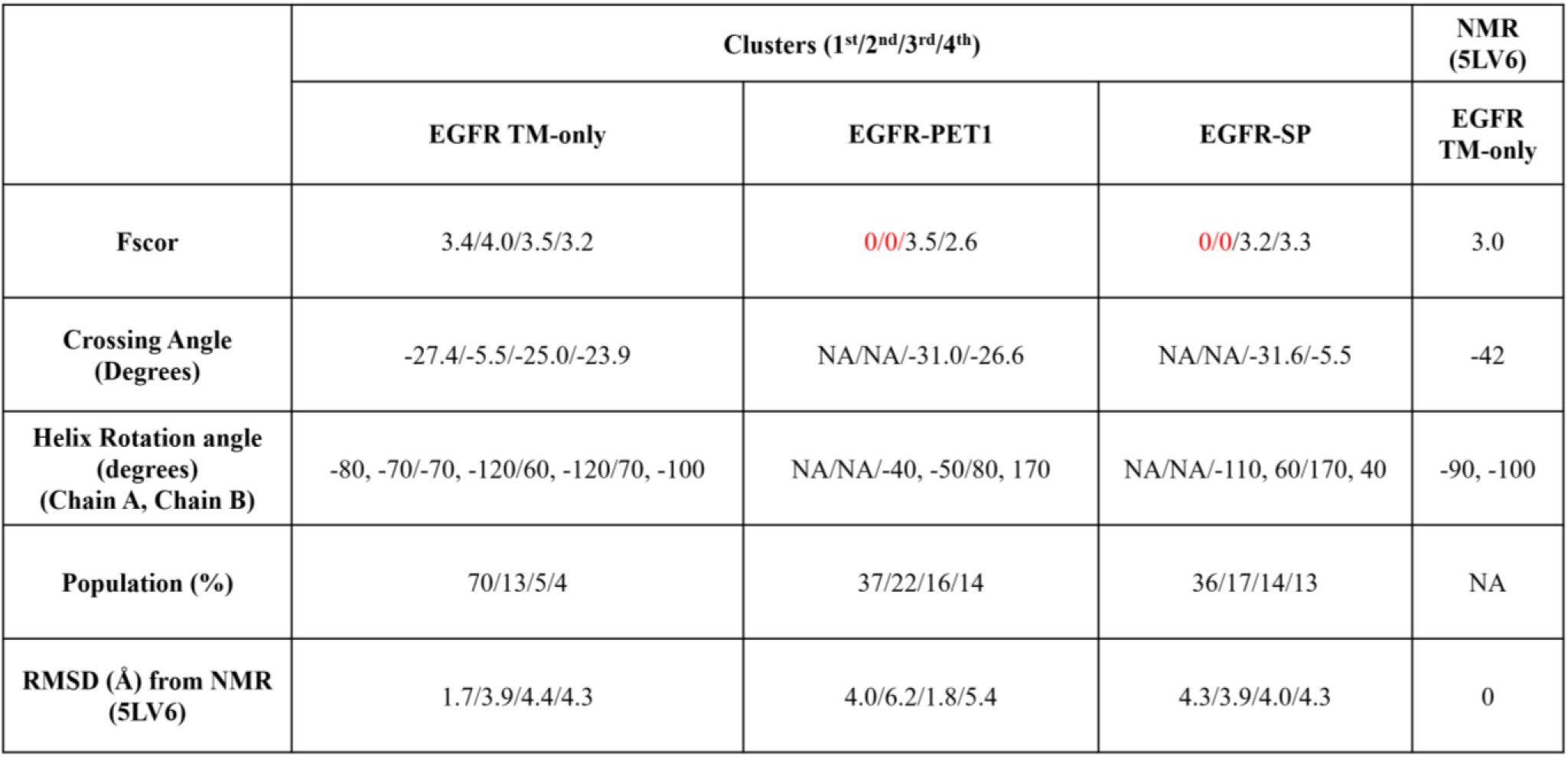
Comparison of CG simulations of EGFR TM-only with EGFR with PET1 and with PET1 scrambled peptide (SP). The last column compares with the NMR EGFR structure. Central conformers of the top 4 clusters for the combined simulations are considered for the comparison using the PREDDIMER method where it compares the Fscor and the crossing angle between the TM helices. Fscor values > 2.5 were considered stable TM dimer structure. In presence of the PET1 peptide, the association of the EGFR TMs are weaker with the Fscor value < 2.5 (shown in red) for the two most populated clusters. PREDDIMER does not calculate the crossing angle if the Fscor is ‘0’ which are the cases for some of the EGFR-PET1 conformers.

|  | Clusters (1<sup>st</sup>/2<sup>nd</sup>/3<sup>rd</sup>/4<sup>th</sup>) |  |  | NMR (5LV6) |
| --- | --- | --- | --- | --- |
|  | EGFR TM-only | EGFR-PET1 | EGFR-SP | EGFR TM-only |
| **Fscor** | 3.4/4.0/3.5/3.2 | 0/0/3.5/2.6 | 0/0/3.2/3.3 | 3.0 |
| **Crossing Angle (Degrees)** | -27.4/-5.5/-25.0/-23.9 | NA/NA/-31.0/-26.6 | NA/NA/-31.6/-5.5 | -42 |
| **Helix Rotation angle (degrees) (Chain A, Chain B)** | -80, -70/-70, -120/60, -120/70, -100 | NA/NA/-40, -50/80, 170 | NA/NA/-110, 60/170, 40 | -90, -100 |
| **Population (%)** | 70/13/5/4 | 37/22/16/14 | 36/17/14/13 | NA |
| **RMSD (Å) from NMR (5LV6)** | 1.7/3.9/4.4/4.3 | 4.0/6.2/1.8/5.4 | 4.3/3.9/4.0/4.3 | 0 |

## Bibliography

1. Wee, P., and Wang, Z. (2017) Epidermal growth factor receptor cell proliferation signaling pathways. Cancers (Basel). 9, 1–45

2. Zeng, F., and Harris, R. C. (2014) Epidermal growth factor, from gene organization to bedside. Semin Cell Dev Biol. 28, 2–11

3. Singh, B., Carpenter, G., and Coffey, R. J. (2016) EGF receptor ligands: Recent advances. F1000Res. 5, 1–11

4. Bocharov, E. v., Lesovoy, D. M., Pavlov, K. v., Pustovalova, Y. E., Bocharova, O. v., and Arseniev, A. S. (2016) Alternative packing of EGFR transmembrane domain suggests that protein-lipid interactions underlie signal conduction across membrane. Biochim Biophys Acta Biomembr. 1858, 1254–1261

5. Bocharov, E. v., Bragin, P. E., Pavlov, K. v., Bocharova, O. v., Mineev, K. S., Polyansky, A. A., Volynsky, P. E., Efremov, R. G., and Arseniev, A. S. (2017) The Conformation of the Epidermal Growth Factor Receptor Transmembrane Domain Dimer Dynamically Adapts to the Local Membrane Environment. Biochemistry. 56, 1697–1705

6. Martin-Fernandez, M. L., Clarke, D. T., Roberts, S. K., Zanetti-Domingues, L. C., and Gervasio, F. L. (2019) Structure and Dynamics of the EGF Receptor as Revealed by Experiments and Simulations and Its Relevance to Non-Small Cell Lung Cancer. Cells. 8, 316

7. Ferguson, K. M. (2008) Structure-based view of epidermal growth factor receptor regulation. Annu Rev Biophys. 37, 353–373

8. Herbst, R. S., and Langer, C. J. (2002) Epidermal growth factor receptors as a target for cancer treatment: The emerging role of IMC-C225 in the treatment of lung and head and neck cancers. Semin Oncol. 29, 27–36

9. Yatabe, Y., and Mitsudomi, T. (2007) Epidermal growth factor receptor mutations in lung cancers. Pathol Int. 57, 233–244

10. Gazdar, A. F. (2009) Activating and resistance mutations of EGFR in non-small-cell lung cancer: Role in clinical response to EGFR tyrosine kinase inhibitors. Oncogene. 28, S24– S31

11. Sabbah, D. A., Hajjo, R., and Sweidan, K. (2020) Review on Epidermal Growth Factor Receptor (EGFR) Structure, Signaling Pathways, Interactions, and Recent Updates of EGFR Inhibitors. Curr Top Med Chem. 20, 815–834

12. Craven, R. J., Lightfoot, H., and Cance, W. G. (2003) A decade of tyrosine kinases: From gene discovery to therapeutics. Surg Oncol. 12, 39–49

13. Cai, W. Q., Zeng, L. S., Wang, L. F., Wang, Y. Y., Cheng, J. T., Zhang, Y., Han, Z. W., Zhou, Y., Huang, S. L., Wang, X. W., Peng, X. C., Xiang, Y., Ma, Z., Cui, S. Z., and Xin, H. W. (2020) The Latest Battles Between EGFR Monoclonal Antibodies and Resistant Tumor Cells. Front Oncol. 10.3389/fonc.2020.01249

14. Huang, L., Jiang, S., and Shi, Y. (2020) Tyrosine kinase inhibitors for solid tumors in the past 20 years (2001–2020). J Hematol Oncol. 13, 1–23

15. Alves, D. S., Westerfield, J. M., Shi, X., Nguyen, V. P., Stefanski, K. M., Booth, K. R., Kim, S., Morrell-Falvey, J., Wang, B.-C., Abel, S. M., Smith, A. W., and Barrera, F. N. (2018) A novel pH-dependent membrane peptide that binds to EphA2 and inhibits cell migration. Elife. 10.7554/eLife.36645

16. Andreev, O. A., Engelman, D. M., and Reshetnyak, Y. K. (2010) PH-sensitive membrane peptides (pHLIPs) as a novel class of delivery agents. Mol Membr Biol. 27, 341–352

17. Nguyen, V. P., Alves, D. S., Scott, H. L., Davis, F. L., and Barrera, F. N. (2015) A Novel Soluble Peptide with pH-Responsive Membrane Insertion. Biochemistry. 54, 6567–6575

18. Nguyen, V. P., Palanikumar, L., Kennel, S. J., Alves, D. S., Ye, Y., Wall, J. S., Magzoub, M., and Barrera, F. N. (2019) Mechanistic insights into the pH-dependent membrane peptide ATRAM. Journal of Controlled Release. 298, 142–153

19. Wei, Y., Liao, R., Mahmood, A. A., Xu, H., and Zhou, Q. (2017) pH-responsive pHLIP (pH low insertion peptide) nanoclusters of superparamagnetic iron oxide nanoparticles as a tumor-selective MRI contrast agent. Acta Biomater. 55, 194–203

20. Estrella, V., Chen, T., Lloyd, M., Wojtkowiak, J., Cornnell, H. H., Ibrahim-Hashim, A., Bailey, K., Balagurunathan, Y., Rothberg, J. M., Sloane, B. F., Johnson, J., Gatenby, R. A., and Gillies, R. J. (2013) Acidity generated by the tumor microenvironment drives local invasion. Cancer Res. 73, 1524–1535

21. Tannock, I. F., and Rotin, D. (1989) Acid pH in tumors and its potential for therapeutic exploitation. Cancer Res. 49, 4373–4384

22. Gatenby, R. A., and Gillies, R. J. (2004) Why do cancers have high aerobic glycolysis? Nat Rev Cancer. 4, 891–899

23. Westerfield, J. M., Sahoo, A. R., Alves, D. S., Grau, B., Cameron, A., Maxwell, M., Schuster, J. A., Souza, P. C. T., Mingarro, I., Buck, M., and Barrera, F. N. (2021) Conformational Clamping by a Membrane Ligand Activates the EphA2 Receptor. J Mol Biol. 433, 167144

24. Scott, H. L., Westerfield, J. M., and Barrera, F. N. (2017) Determination of the Membrane Translocation pK of the pH-Low Insertion Peptide. Biophys J. 113, 869–879

25. Mineev, K. S., Bocharov, E. V, Pustovalova, Y. E., Bocharova, O. V, Chupin, V. V, and Arseniev, A. S. (2010) Spatial structure of the transmembrane domain heterodimer of ErbB1 and ErbB2 receptor tyrosine kinases. J Mol Biol. 400, 231–243

26. Endres, N. F., Das, R., Smith, A. W., Arkhipov, A., Kovacs, E., Huang, Y., Pelton, J. G., Shan, Y., Shaw, D. E., Wemmer, D. E., Groves, J. T., and Kuriyan, J. (2013) Conformational coupling across the plasma membrane in activation of the EGF receptor. Cell. 152, 543–556

27. Chattopadhyay, A. (1990) Chemistry and biology of N-(7-nitrobenz-2-oxa-1,3-diazol-4-yl)-labeled lipids: fluorescent probes of biological and model membranes. Chem Phys Lipids. 53, 1–15

28. Johnson, A. E. (2005) Fluorescence approaches for determining protein conformations, interactions and mechanisms at membranes. Traffic. 6, 1078–1092

29. Mazères, S., Schram, V., Tocanne, J. F., and Lopez, A. (1996) 7-nitrobenz-2-oxa-1,3-diazole-4-yl-labeled phospholipids in lipid membranes: differences in fluorescence behavior. Biophys J. 71, 327–335

30. Vivian, J. T., and Callis, P. R. (2001) Mechanisms of tryptophan fluorescence shifts in proteins. Biophys J. 80, 2093–2109

31. Makowiecka, A., Simiczyjew, A., Nowak, D., and Mazur, A. J. (2016) Varying effects of EGF, HGF and TGFγ on formation of invadopodia and invasiveness of melanoma cell lines of different origin. European Journal of Histochemistry. 60, 230–238

32. Pietraszek-Gremplewicz, K., Simiczyjew, A., Dratkiewicz, E., Podgórska, M., Styczeń, I., Matkowski, R., Ziętek, M., and Nowak, D. (2019) Expression level of EGFR and MET receptors regulates invasiveness of melanoma cells. J Cell Mol Med. 23, 8453–8463

33. Alves, D. S., Westerfield, J. M., Shi, X., Nguyen, V. P., Stefanski, K. M., Booth, K. R., Kim, S., Morrell-Falvey, J., Wang, B.-C., Abel, S. M., Smith, A. W., and Barrera, F. N. (2018) A novel pH-dependent membrane peptide that binds to EphA2 and inhibits cell migration. Elife. 7, e36645

34. Christie, S., Shi, X., and Smith, A. W. (2020) Resolving Membrane Protein-Protein Interactions in Live Cells with Pulsed Interleaved Excitation Fluorescence Cross-Correlation Spectroscopy. Acc Chem Res. 53, 792–799

35. Dunn, K. W., Kamocka, M. M., and McDonald, J. H. (2011) A practical guide to evaluating colocalization in biological microscopy. Am J Physiol Cell Physiol. 300, 723–742

36. Yan, D., Parker, R. E., Wang, X., Frye, S. v., Shelton Earp, H., DeRyckere, D., and Graham, D. K. (2018) MERTK promotes resistance to irreversible EGFR tyrosine kinase inhibitors in non–small cell lung cancers expressing wild-type EGFR family members. Clinical Cancer Research. 24, 6523–6535

37. Liu, W., Yu, C., Li, J., and Fang, J. (2022) The Roles of EphB2 in Cancer. Front Cell Dev Biol. 10.3389/fcell.2022.788587

38. Kennedy, S. P., Hastings, J. F., Han, J. Z. R., and Croucher, D. R. (2016) The under-appreciated promiscuity of the epidermal growth factor receptor family. Front Cell Dev Biol. 10.3389/fcell.2016.00088

39. Evans, R., O’neill, M., Pritzel, A., Antropova, N., Senior, A., Green, T., Žídek, A., Bates, R., Blackwell, S., Yim, J., Ronneberger, O., Bodenstein, S., Zielinski, M., Bridgland, A., Potapenko, A., Cowie, A., Tunyasuvunakool, K., Jain, R., Clancy, E., Kohli, P., Jumper, J., and Hassabis, D. (2022) Protein complex prediction with AlphaFold-Multimer. bioRxiv. 10.1101/2021.10.04.463034

40. Endres, N. F., Das, R., Smith, A. W., Arkhipov, A., Kovacs, E., Huang, Y., Pelton, J. G., Shan, Y., Shaw, D. E., Wemmer, D. E., Groves, J. T., and Kuriyan, J. (2013) Conformational coupling across the plasma membrane in activation of the EGF receptor. Cell. 152, 543–556

41. Kim, D. H., Triet, H. M., and Ryu, S. H. (2021) Regulation of EGFR activation and signaling by lipids on the plasma membrane. Prog Lipid Res. 83, 101115

42. Kovacs, E., Zorn, J. A., Huang, Y., Barros, T., and Kuriyan, J. (2015) A Structural Perspective on the Regulation of the Epidermal Growth Factor Receptor. Annu Rev Biochem. 84, 739–764

43. Arkhipov, A., Shan, Y., Das, R., Endres, N. F., Eastwood, M. P., Wemmer, D. E., Kuriyan, J., and Shaw, D. E. (2013) Architecture and Membrane Interactions of the EGF Receptor. Cell. 152, 557–569

44. Lelimousin, M., Limongelli, V., and Sansom, M. S. P. (2016) Conformational Changes in the Epidermal Growth Factor Receptor: Role of the Transmembrane Domain Investigated by Coarse-Grained MetaDynamics Free Energy Calculations. J Am Chem Soc. 138, 10611–10622

45. Sinclair, J. K. L., Walker, A. S., Doerner, A. E., and Schepartz, A. (2018) Mechanism of Allosteric Coupling into and through the Plasma Membrane by EGFR. Cell Chem Biol. 25, 857–870.e7

46. Huang, Y., Ognjenović, J., Karandur, D., Miller, K., Merk, A., Subramaniam, S., and Kuriyan, J. (2021) A molecular mechanism for the generation of Ligand-dependent differential outputs by the epidermal growth factor receptor. Elife. 10, 1–32

47. Du, Z., Brown, B. P., Kim, S., Ferguson, D., Pavlick, D. C., Jayakumaran, G., Benayed, R., Gallant, J. N., Zhang, Y. K., Yan, Y., Red-Brewer, M., Ali, S. M., Schrock, A. B., Zehir, A., Ladanyi, M., Smith, A. W., Meiler, J., and Lovly, C. M. (2021) Structure–function analysis of oncogenic EGFR Kinase Domain Duplication reveals insights into activation and a potential approach for therapeutic targeting. Nat Commun. 12, 1382

48. Huang, Y., Bharill, S., Karandur, D., Peterson, S. M., Marita, M., Shi, X., Kaliszewski, M. J., Smith, A. W., Isacoff, E. Y., and Kuriyan, J. (2016) Molecular basis for multimerization in the activation of the epidermal growth factor receptor. Elife. 5, e14107

49. Sali, A., and Blundell, T. L. (1993) Comparative protein modelling by satisfaction of spatial restraints. J Mol Biol. 234, 779–815

50. Souza, P. C. T., Alessandri, R., Barnoud, J., Thallmair, S., Faustino, I., Grünewald, F., Patmanidis, I., Abdizadeh, H., Bruininks, B. M. H., Wassenaar, T. A., Kroon, P. C., Melcr, J., Nieto, V., Corradi, V., Khan, H. M., Domański, J., Javanainen, M., Martinez-Seara, H., Reuter, N., Best, R. B., Vattulainen, I., Monticelli, L., Periole, X., Tieleman, D. P., de Vries, A. H., and Marrink, S. J. (2021) Martini 3: a general purpose force field for coarse-grained molecular dynamics. Nat Methods. 18, 382–388

51. Abraham, M. J., Murtola, T., Schulz, R., Páll, S., Smith, J. C., Hess, B., and Lindahl, E. (2015) GROMACS: High performance molecular simulations through multi-level parallelism from laptops to supercomputers. SoftwareX. 1–2, 19–25

52. Wassenaar, T. A., Ingólfsson, H. I., Böckmann, R. A., Tieleman, D. P., and Marrink, S. J. (2015) Computational Lipidomics with insane: A Versatile Tool for Generating Custom Membranes for Molecular Simulations. J Chem Theory Comput. 11, 2144–2155

53. Polyansky, A. A., Chugunov, A. O., Volynsky, P. E., Krylov, N. A., Nolde, D. E., and Efremov, R. G. (2014) PREDDIMER: a web server for prediction of transmembrane helical dimers. Bioinformatics. 30, 889–890

54. Larkin, M. A., Blackshields, G., Brown, N. P., Chenna, R., McGettigan, P. A., McWilliam, H., Valentin, F., Wallace, I. M., Wilm, A., Lopez, R., Thompson, J. D., Gibson, T. J., and Higgins, D. G. (2007) Clustal W and Clustal X version 2.0. Bioinformatics. 23, 2947–2948

